# The genetic landscape of antibiotic sensitivity in Staphylococcus aureus

**DOI:** 10.1101/2024.08.15.608136

**Authors:** Wan Li, Menghan Liu, Panos Oikonomou, Sydney Blattman, Falguni Paul, Julia Hettleman, Joseph Gonzalez, Qiaoyu Hu, Howe Chen, Saeed Tavazoie, Wenyan Jiang

## Abstract

A comprehensive genetic landscape of antibiotic sensitivity in *Staphylococcus aureus* is lacking. Using ultra-dense CRISPR-interference libraries, we systematically quantified global gene fitness across ten antibiotics and uncovered hundreds of significant antibiotic-gene interactions. Essential genes dominated these interactions, a finding not revealed by transposon-based studies. Processes most vulnerable to transcriptional repression under antibiotic conditions included cell wall synthesis/cell division (CC), DNA replication/DNA recombination (DD), coenzyme A biosynthesis, and riboflavin metabolism. Network and genetic analyses further revealed novel synergistic genetic interactions (GIs) within these processes, including an extensive CC-DD subnetwork. Only a subset of CC-DD synergies was dependent on the cell division inhibitor SosA. Informed by these GIs, we identified multiple drug-drug combinations with potent synergistic activity against multidrug-resistant *S. aureus*. Our detailed profiling of drug-gene, gene-gene, and drug-drug interactions reveals novel functional relationships among essential genes and defines a vulnerability landscape to guide new drug target discovery and effective combination therapies.

## INTRODUCTION

The rise of antimicrobial resistance (AMR) is one of the most pressing threats to global public health. In 2021, bacterial AMR was associated with 4.71 million deaths worldwide^1^. Without effective intervention, AMR is projected to surpass cancer as the leading cause of death by 2050, with an estimated annual economic burden of US $3.5 billion^2^. Among bacterial pathogens, *Staphylococcus aureus* stands out for its remarkable ability to withstand antibiotics and evade the human immune system, making it a leading cause of infection-related morbidity and mortality^1^. *S. aureus* possesses many intrinsic factors that limit antibiotic efficacy. Previous studies using transposon insertion sequencing (Tn-seq) have allowed researchers to survey the contribution of every nonessential gene in the *S. aureus* core genome to bacterial fitness under antibiotic stress^3,4^. These studies have shown that perturbation of diverse biological processes, including cell wall and membrane function (eg, *mprF*, *tagO*), electron transport chain (eg, *qoxAB*), and two-component systems (eg, *graRS*, *vraFG*) can strongly modulate antimicrobial sensitivity.

Transposon insertion techniques exert strong, irreversible perturbations, limiting their utility for studying phenotypes involving essential genes. Consequently, systems-level investigations of how essential genes influence antibiotic sensitivity in *S. aureus* have been lacking. Yet, this knowledge is critical both for advancing our understanding of microbial genetic architecture and for guiding new antimicrobial strategies. Indeed, systems biology studies have shown that essential genes occupy central positions in genetic interaction (GI) networks in yeast^5^ and exert markedly stronger effects on antibiotic sensitivity than nonessential genes in *Mycobacterium tuberculosis*^6^. Insights from these studies could inform the development of effective combinatorial drug therapies, which have proven successful across many medical fields, including infectious diseases^7-9^, HIV^10^, cancer^11^, and cardiovascular disease^12^. Compared to monotherapies, combinatorial regimens offer multiple advantages, including synergistic efficacy and resistance repression, which can reduce toxicity within human hosts and improve overall therapeutic outcomes.

To investigate the effects of essential genes on bacterial phenotypes, we previously developed CRISPR interference (CRISPRi) technology^13,14^, which mediates mild transcriptional repression rather than strong gene disruption. Recent applications of CRISPRi in bacteria by us and others have already revealed new roles essential genes play in antibiotic sensitivity^6,15-18^, showing that the genetic landscapes profiled by Tn-seq are incomplete. For instance, repression of the essential two-component system MtrAB sensitizes *M. tuberculosis* to multiple antibiotics, including rifampicin, vancomycin, and linezolid^6^. Using single- and dual-CRISPRi libraries in *S. aureus*, we showed that repression of the essential mevalonate and fatty acid synthesis pathways enhances and diminishes, respectively, intrinsic resistance to gentamicin^15^.

In this study, we applied a large-scale systems genomics approach to map the antibiotic sensitivity landscape of *S. aureus* across ten antibiotics that target cell wall synthesis/membrane, DNA synthesis/replication, and protein translation. Leveraging CRISPR Adaptation-mediated Library Manufacturing (CALM), a CRISPRi functional genomics platform we developed previously^15^, we generated highly comprehensive genome-scale libraries in both methicillin-susceptible *S. aureus* (MSSA) and methicillin-resistant *S. aureus* (MRSA). Our screens revealed hundreds of significant antibiotic-gene interactions, with essential genes dominating these interactions. Repression of essential genes could either sensitize or desensitize *S. aureus* to antibiotics, a finding not revealed by previous transposon-based studies^3,4^. Among the most vulnerable biological processes were <u>c</u>ell wall synthesis/<u>c</u>ell division (CC), <u>D</u>NA replication/<u>D</u>NA recombination (DD), coenzyme A biosynthesis, and riboflavin metabolism, as their repression significantly reduced bacterial fitness in antibiotics of various modes of action. The scale of our antibiotic-gene interaction dataset enabled the construction of an essential gene similarity network, which uncovered novel synergistic GIs among essential genes within these processes. Finally, we translated several newly identified synergistic GIs into antimicrobial combinations that exhibited potent killing against both MSSA and multidrug-resistant MRSA. Together, our comprehensive profiling of drug-gene, gene-gene, and drug-drug interactions illuminates new functional relationships within the *S. aureus* core essential genome and provides a rational foundation for developing effective combinatorial antimicrobial therapies. Our work also places *S. aureus* among the few pathogens for which genome-scale CRISPRi-based characterization of antibiotic sensitivity has been achieved, alongside *M. tuberculosis*^6^.

## RESULTS

Harnessing the natural capacity of CRISPR adaptation – a bacterial immunization process that stores nucleic acid sequence information from invaders such as phages – the CALM method^15^ can rapidly generate comprehensive genome-scale CRISPR RNA (crRNA) libraries that cover 95% of all targetable sites in the *S. aureus* genome, bypassing the conventional, resource-intensive steps of guide RNA (gRNA) synthesis and cloning. As outlined in **Fig. 1a**, *S. aureus* cells harboring an engineered CRISPR adaptation machinery convert genomic DNA into ultra-dense genome-scale crRNA libraries. The canonical crRNA, along with another small RNA encoded by *tracr*, is functionally equivalent to the engineered gRNA^19^. Both crRNA and dCas9 are constitutively expressed. To profile the genetic landscape of antibiotic sensitivity, libraries generated in strains RN4220^20^ and JE2^21^ (a MRSA USA300 strain^22^ cured of all its plasmids) were cultured in plain tryptic soy broth (TSB) media or TSB containing antibiotics targeting cell wall synthesis/membrane, DNA synthesis/replication, and protein translation (**Fig. 1b**). The ten antibiotics were supplied at sub-lethal concentrations (**Fig. S1a**), and bacterial cultures were grown for 9 hours, reaching early stationary phase.

**Fig. 1.**
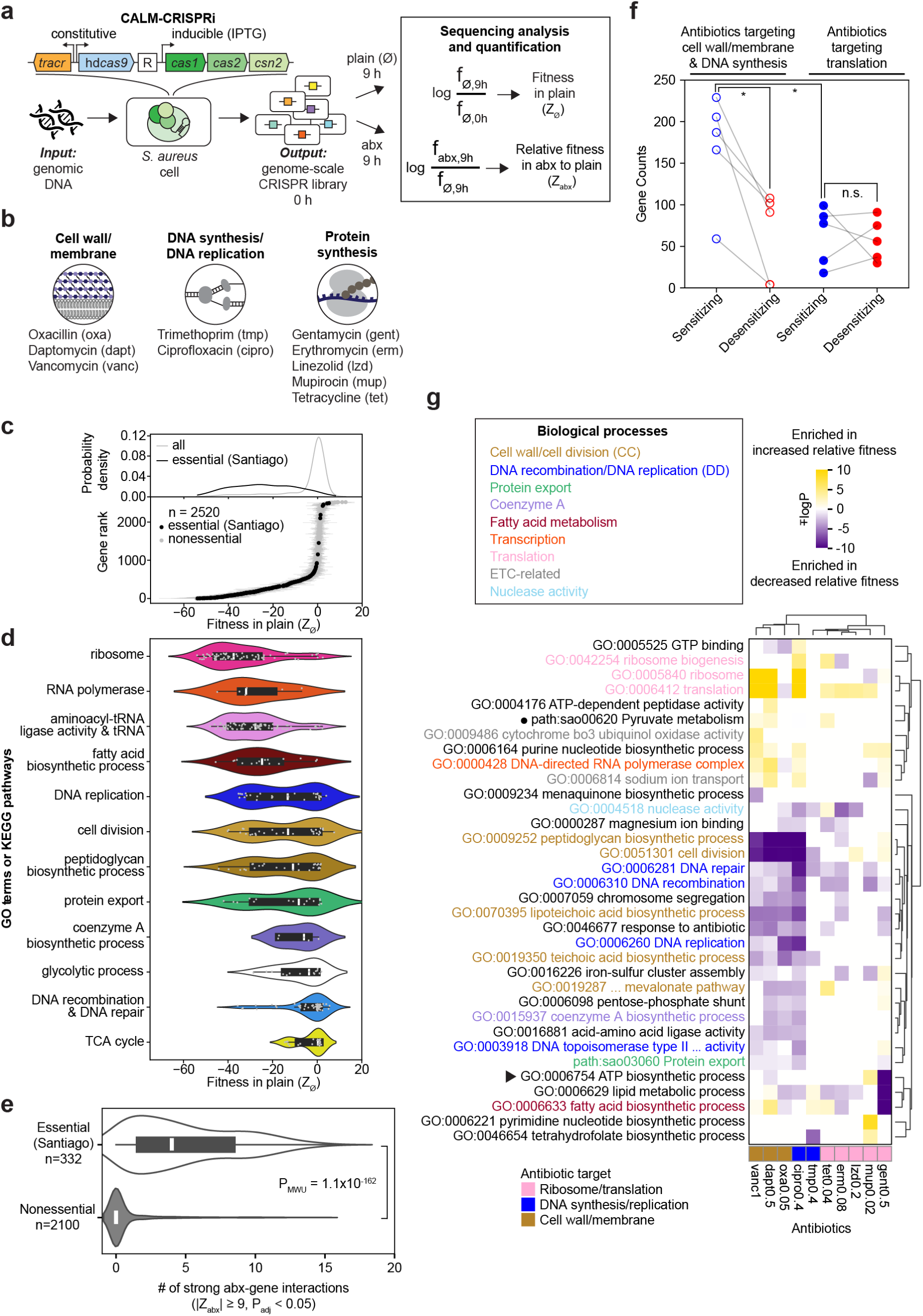
CALM-CRISPRi reveals the genetic landscape of essentiality and antibiotic sensitivity in *Staphylococcus aureus*. (**a**) Utilizing CRISPR Adaptation-mediated Library Manufacturing (CALM) technology, highly comprehensive genome-scale CRISPRi libraries can be generated in *S. aureus*. *tracr*, hd*Cas9* (hyper dCas9) and R (crRNA), which are components necessary for transcriptional repression, are constitutively expressed. These libraries were subjected to outgrowth in plain TSB (Ø) or sublethal concentrations of ten antibiotics (abx), followed by sequencing analysis and quantification. (**b**) The ten antibiotics used in current study. (**c**) Distribution of gene fitness in plain TSB (Z_Ø_). Lower panel: ranked *S. aureus* RN4220 genes based on their mean fitness in TSB (Z_Ø_) quantified by CRISPRi from three biological replicates, with error bars showing the standard deviations. Essential and nonessential genes as determined by Santiago’s Tn-seq study^23^ are shown as black and gray dots, respectively. Upper panel: distributions of Santiago’s essential genes (black) and all genes (gray). Z_Ø_ values are shown in Table S2. (**d**) Distributions of Z_Ø_ for core essential biological processes. (**e**) Antibiotic-gene interactions by gene essentiality. Violin plot showing the distributions of the number of antibiotic conditions in which repression of essential and nonessential genes significantly altered relative fitness (|Z_abx_| ≥ 9 and P_adj_ < 0.05). A Mann-Whitney U test was performed for the two gene groups. (**f**) The numbers of significant sensitizing (blue) and desensitizing (red) antibiotic-gene interactions (|Z_abx_| ≥ 9 and P_adj_ < 0.05) for different antibiotic groups. Wilcoxon test and Mann-Whitney U test were performed for paired and non-paired groups, respectively. * indicates P < 0.05. (**g**) Functional enrichment analysis by iPAGE, an information-theoretic pathway analysis tool^60^, followed by hierarchical clustering of significantly enriched non-redundant Gene Ontology (GO) terms and KEGG pathways. The ∓log(P-value) of GO terms and KEGG pathways significantly enriched (P < 0.001) for genes whose repression increased and decreased relative fitness in antibiotics (|Z_abx_| ≥ 9 and P_adj_ < 0.05) are shown in yellow and purple, respectively. Clustering of all GO terms and KEGG pathways are shown in Figs. S4a, S4c, Tables S6, and S8.

The CRISPRi libraries generated by CALM were highly diverse, with nearly 90% of genes covered by 3 or more coding-strand-targeting (CT) crRNAs and a median of ≥10 CT crRNAs per gene (**Fig. S1b**), conferring statistical robustness. We used CT crRNAs to quantify gene fitness, as they mediate strong transcriptional repression^13,14^. For each crRNA, we quantified its fitness in plain media 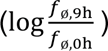 and the relative fitness in antibiotic compared to plain media 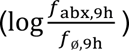 (**Fig. 1a**). Next, a gene score was calculated by averaging the values from all CT crRNAs targeting that gene. Finally, we standardized the gene score by using a pool of null, intergenic crRNAs, yielding a Z-score (Z_Ø_ and Z_abx_) as the final fitness score (**Fig. 1a** and Methods). This standardization allows direct comparison of gene fitness effects across different samples and antibiotic treatments – for example, a Z-score of ±1 indicates that the fitness effect of a gene’s repression is one standard deviation from the mean of the null distribution. Since Z_abx_ reflects the relative fitness in antibiotic compared to plain media, negative and positive Z_abx_ values indicate sensitization and desensitization of bacteria to antibiotics, respectively. Z-scores from independent biological replicates showed good Pearson correlations: 0.89 < r < 0.97 in plain media (**Figs. S1d, S1e**) and 0.77 < r < 0.93 in antibiotic conditions (**Fig. S1f**).

Our CALM-CRISPR libraries showed good performance in identifying essential genes: 0.91 < AUC < 0.95 (**Figs. S1g, S1h**); genes quantified with low fitness (Z_Ø_) in plain media significantly overlapped with essential genes identified in Santiago’s Tn-seq study^23^ (**Fig. 1c**). The degree of essentiality quantified by CRISPRi varied widely across core essential biological processes, ranging from highly essential genes encoding the ribosome, RNA polymerase, tRNA ligase/tRNA, and fatty acid biosynthesis pathway, to less essential genes involved in coenzyme A biosynthesis, glycolysis, DNA recombination/DNA repair, and TCA cycle (**Figs. 1d** and **S2**). A similar gene differential essentiality profile was also found in Mycobacterium^24^, suggesting that gene vulnerability may be at least partially conserved across microbial species.

### A genetic landscape of antibiotic sensitivity dominated by essential genes

650 out of ∼2600 annotated genes in the RN4220 genome showed significantly altered relative fitness (|Z_abx_| ≥ 9 and P_adj_ < 0.05) in at least one antibiotic condition compared to plain media, among a total of 18 conditions we tested (**Fig. S3a**). Hierarchical clustering of these genes formed two major groups. The lower group consisted of antibiotics that target cell wall/membrane and DNA replication, which are vancomycin, daptomycin, oxacillin, and ciprofloxacin. The upper group included mostly antibiotics targeting protein synthesis, with trimethoprim as an exception. Notably, we found that essential genes dominated the antibiotic-gene interactions. Perturbation of 66% of essential genes identified by Santiago’s^23^ and Bae’s^25^ Tn-seq studies significantly altered relative fitness in antibiotics (black bars in **Fig. S3a**). Overall, perturbation of essential genes led to strong changes in relative fitness (|Z_abx_| ≥ 9 and P_adj_ < 0.05) in a median of 4 antibiotic conditions, while perturbation of ∼80% nonessential genes did not significantly alter relative fitness in any antibiotic condition (**Fig. 1e**). These results show that CRISPRi, with its ability to mildly perturb essential genes, revealed a new range of antibiotic-gene interactions not attainable with the strong loss-of function Tn-seq methodology. Only 1.7% - 7.1% of nonessential genes were found to significantly modulate relative fitness in antibiotic conditions (**Fig. S3b**), and they were enriched in processes such as ATP biosynthesis and electron transport chain (ETC) (**Fig. S4d**). Additionally, among the interactions between genes and antibiotics that target cell wall/membrane and DNA synthesis, sensitizing interactions significantly outnumbered desensitizing interactions (P_Wilcoxon_ < 0.05) (**Fig. 1f**), indicating that gene repression more frequently decreased, rather than increased, relative fitness in these antibiotics. Sensitizing interactions were also significantly more prevalent for these antibiotics than for those targeting translation (P_Mann-Whitney U_ < 0.05) (**Fig. 1f**).

To gain insights into the biological functions of genes that modulate bacterial fitness in the antibiotics, we performed functional enrichment analysis and clustering. This analysis identified significantly overrepresented Gene Ontology (GO) terms (**Figs. 1g** and **S4a,**) and KEGG pathways (**Fig. S4c**) among genes whose repression decreased (purple) or increased relative fitness (yellow) in antibiotics. As expected, inhibition of many biological processes sensitized bacteria to antibiotics with related modes of action. For instance, CC processes (brown labels in **Fig. 1g**) were highly enriched among genes whose repression reduced relative fitness in cell-wall/membrane-targeting antibiotics such as oxacillin, daptomycin, and vancomycin. Likewise, DD processes (blue labels in **Fig. 1g**) were enriched among genes whose repression reduced relative fitness in ciprofloxacin, which target DNA replication. Because bacterial genes are organized in operons, functional enrichment can be confounded as a result of silencing nearby co-transcribed genes involved in different biological processes. To address this, we also performed a more conservative enrichment analysis using only the last gene in each operon (**Table S1**). Although removing ∼1,000 genes reduced the sensitivity of the analysis, this last-gene-in-operon approach still produced consistent enrichment profiles, with antibiotic conditions showing an average Pearson correlation of 0.80 compared to profiles generated using all genes (**Fig. S4b**).

### Repression of CC and DD processes sensitize bacteria to many antibiotics

Our CALM screens revealed that CC processes were among the most vulnerable, as their knockdown sensitized bacteria to many antibiotics we tested (**Figs. 1g** and **S4a**). In addition to the antibiotics that target cell wall/membrane, many CC processes were also significantly enriched among genes whose repression reduced relative fitness in DNA synthesis/replication-targeting antibiotics (ie, trimethoprim and ciprofloxacin). At the gene level, we found that inhibition of a substantial portion of essential genes in peptidoglycan biosynthesis (GO:0009252) and cell division (GO:0051301) significantly decreased relative fitness in ciprofloxacin and trimethoprim (**Figs. 2a-2d**), a phenotype not captured by a previous Tn-study^3^. To validate these CRISPRi screening results, we cloned CT crRNAs targeting representative essential gene hits, *ftsZ* and *femX*, with an IPTG-inducible dCas9 and quantified their fitness relative to that of a non-targeting control in antibiotics (Pairwise competition assay in Methods). Whenever possible, we chose to validate genes that are the last in operons (**Fig. S5a**) to avoid polar effects of CRISPRi. Importantly, we used IPTG concentrations that moderately reduced the fitness of candidate cells to 50-90% of that of the non-targeting control under no antibiotic treatment, as excessive inhibition of essential genes led to strong growth defect (or death) even without antibiotics (**Fig. S5a**). Our approach confirmed that inhibition of *ftsZ* and *femX* significantly reduced relative fitness in oxacillin, daptomycin, vancomycin, ciprofloxacin, and trimethoprim (**Fig. 2e**). Similarly, inhibition of other essential processes upstream of peptidoglycan synthesis, such as the mevalonate pathway^26^ (**Figs. S4e** and **S4f**) that synthesizes the precursor to undecaprenyl phosphate, and pathways encoding cell surface glycopolymers specific to Gram-positive bacteria, including teichoic acid and lipoteichoic acid biosynthetic processes^27^, also sensitized bacteria to these five antibiotics (**Figs. S4g-S4i**).

**Fig. 2.**
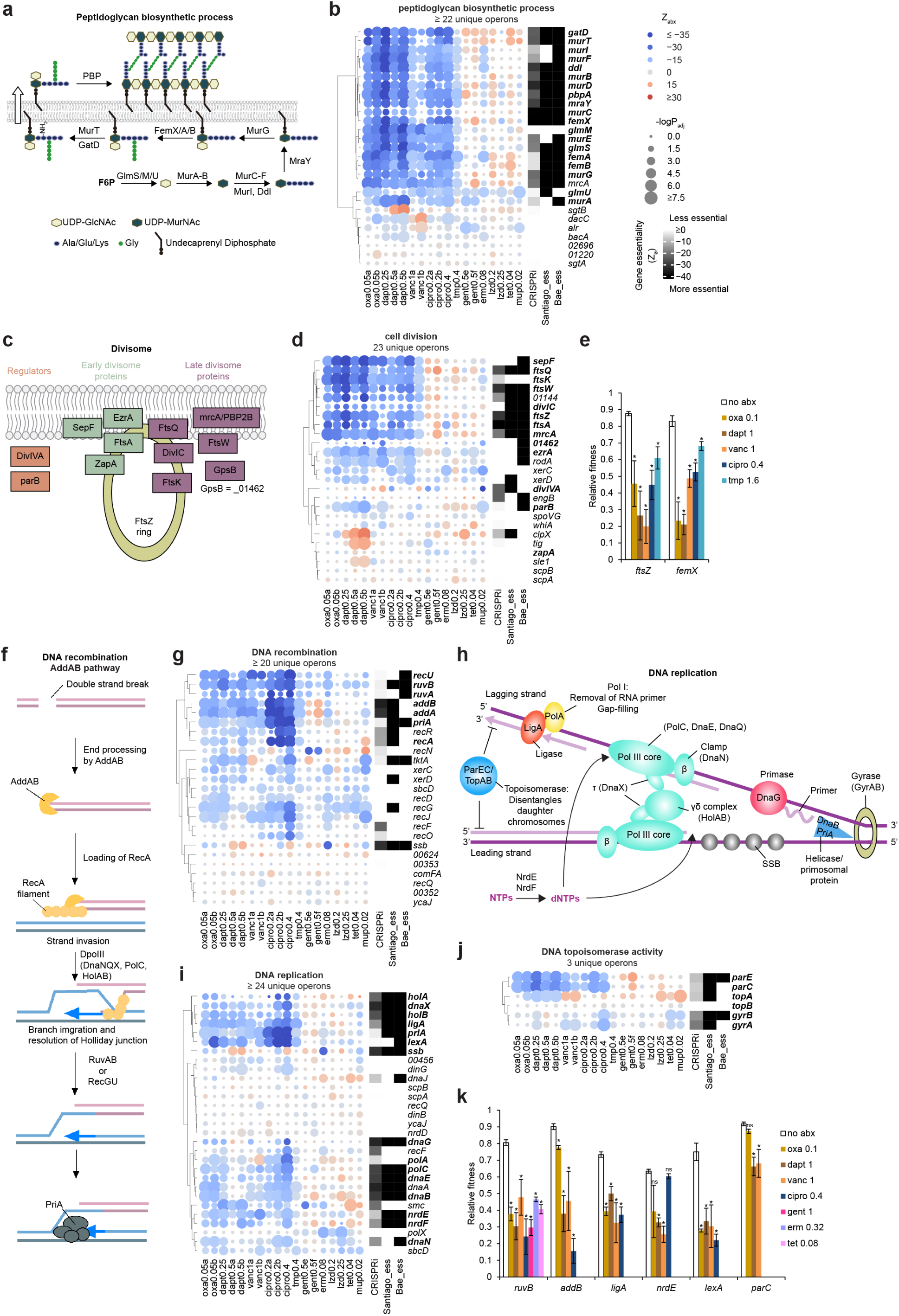
Repression of CC and DD processes sensitize bacteria to many antibiotics. (**a**) Peptidoglycan biosynthetic pathway. (**b**) Heatmap showing the relative fitness of genes in the peptidoglycan biosynthetic process (GO:0009252) in ten antibiotics as quantified by CALM-CRISPRi screens (Z_abx_) in *S. aureus* RN4220. Gene names are annotated by KEGG orthology (ko). *glmS* and *pbpA* were also included due to overannotation by GO terms. Genes that appear in schematic in (a) are bolded. Gene knockdowns that decreased and increased relative fitness in antibiotics are shown in blue and red circles, respectively. Size of circle indicates the negative of log-transformed adjusted P-value, - logP_adj_. Gene essentiality quantified by CRISPRi (mean of Z_Ø_ from triplicates) and qualified (binary) by Santiago’s^23^ and Bae’s^25^ Tn-seq studies are also shown. (**c**) Schematic of the divisome. (**d**) Same as (b) except heatmap is showing genes in cell division (GO:0051301). *ftsK*, *mrcA*, and *zapA* were also included due to overannotation by GO terms. *mur* genes already shown in (b) were omitted. (**e**) Pairwise competition assays (Methods) measuring the relative fitness of *S. aureus* RN4220 carrying an IPTG-inducible CRISPRi system targeting *ftsZ* and *femX* in indicated antibiotic conditions. Concentrations of antibiotics are shown in µg/mL. Error bars indicate the standard deviation from three biological replicates. For each gene knockdown, paired t-tests were performed between the antibiotic condition and the no-antibiotic condition. * indicates P < 0.05. (**f**) The AddAB (Gram-positive homologue of RecBCD) DNA recombination pathway. Schematic is adapted from KEGG (www.genome.jp/kegg/) with modifications. (**g**) Same as (b) except heatmap is showing genes in DNA recombination (GO:0006310). *addA*, *addB* and *recF* were also included due to overannotation by GO terms. (**h**) Schematic of DNA replication. (**i, j**) Same as (b) except heatmaps are showing genes in DNA replication (GO:0006260) and DNA topoisomerase activity (GO:0003916), respectively. (**k**) Same as (e) except bar plots are showing the relative fitness of repressions of *ruvB*, *addB*, *ligA*, *nrdE*, *lexA*, and *parC*.

DD processes were also highly vulnerable, as their repression sensitized bacteria to many antibiotics (**Figs. 1g** and **S4a**). As expected, knockdown of genes encoding the two bacterial DNA homologous recombination (GO:0006310) pathways, RecFOR and AddAB (the RecBCD homologue in *S. aureus*^28^, **Fig. 2f**), led to the greatest reduction in relative fitness in ciprofloxacin (genes bolded in **Fig. 2g**). Relative fitness in other antibiotics, especially those that target cell wall/membrane also significantly decreased. For example, Z_abx_ values of *addAB*, encoding the Gram-positive RecBCD homologue, and those of *ruvAB*, encoding the essential Holliday junction DNA helicase, were all around -15 in oxacillin, daptomycin, and vancomycin. Repression of additional essential DD genes (**Figs. 2h-2j**) – including those encoding the DNA replication machinery (eg, *ligA*, *polC*, *dnaEX*, *holAB*, *priA*, *dnaB*), the ribonucleoside-diphosphate reductase (*nrdEF*) that converts NTPs to dNTPs, the DNA repair regulator *lexA*, and DNA topoisomerase IV (*parEC*) – also reduced relative fitness in all or a subset of the three cell wall/membrane-targeting antibiotics. Using pairwise competition, we validated the reduced relative fitness of *ruvB*, *addB*, *ligA*, *nrdE*, *lexA* and *parC* in most antibiotic conditions (**Fig. 2k**), with three conditions in *nrdE* and *parC* not reaching statistical significance. Notably, *ruvB* knockdown also sensitized bacteria to translation inhibitors including gentamicin, erythromycin, and tetracycline (**Figs. 2g** and **2k**). Supporting these results obtained with pairwise competition, repression of these DD genes, except for *parC*, showed a clear reduction in their minimum inhibitory concentration (MIC) in oxacillin (**Figs. S5h, S5i** and **S5n-S5q**). Since *ruvB* and *ligA* precede genes of other functions in their respective operons (**Fig. S5a**), we also conducted complementation experiments to confirm their roles in antibiotic sensitivity (**Figs. S5r-S5u)**.

We also performed CALM-CRISPRi screens in a MRSA strain, JE2, and found that repression of many CC genes (eg, *glmM*, *murF*, *ftsZ*, and *sepF* in **Figs. S6a** and **S6b**) and DD genes (eg, *ruvB*, *nrdE*, *priA*, *polC*, *lexA*, and *parC* in **Figs. S6c-S6e**) significantly reduced relative fitness in daptomycin and vancomycin, antibiotics commonly used for treating MRSA infection. Together, these findings revealed CC and DD as highly vulnerable processes where their inhibition sensitizes *S. aureus* to various antibiotics, especially those that target cell wall/membrane and DNA synthesis.

### Other prominent sensitizing antibiotic-gene interactions

Our functional enrichment analysis identified additional essential pathways whose inhibition strongly sensitized bacteria to antibiotics that target cell wall/membrane and DNA synthesis. Many of these pathways, including protein export (green label in **Fig. 1g**) and the biosynthesis of essential organic cofactors such as coenzyme A (purple label in **Fig. 1g**), flavin adenine dinucleotide (FAD), and nicotinamide adenine dinucleotide (NAD), were not captured by previous transposon studies^3,4^. Protein integration and transport across the membranes is an essential process ubiquitous to all domains of life. In bacteria, transmembrane proteins are inserted into the membrane by the signal recognition particle (SRP) and the Sec translocation machinery and channels^29^ (**Fig. 3a**). We found that inhibition of a subset of protein transport components, especially the core of SRP (4.5S RNA and Ffh) and the ATPase motor of the Sec translocation machinery (SecA), potentiated oxacillin, daptomycin, vancomycin, and ciprofloxacin (**Figs. 3b** and **3c**). Inhibition of biosynthetic processes of a number of essential organic cofactors central to cellular metabolism, such as coenzyme A, FAD, and NAD, also potentiated these four antibiotics (**Figs. 3d-3g, S4j**, and **S4k**). We confirmed that knockdown of the enzymes responsible for producing these cofactors, including the essential *coaA*, *ribF*, and *ppnK* indeed reduced relative fitness in these antibiotics (**Fig. 3h**), as well as the MICs of oxacillin (**Figs. S5e, S5k** and **S5l**) and ciprofloxacin (**Fig. S5v**). For bacteria with *ribF* knockdown, we also chemically rescued the phenotype by supplementing growth media with FAD (**Fig. 3h**), a key cofactor synthesized by RibF (**Fig. 3f**). This indicates that lack of FAD sensitizes bacteria to various antibiotic stress.

**Fig. 3.**
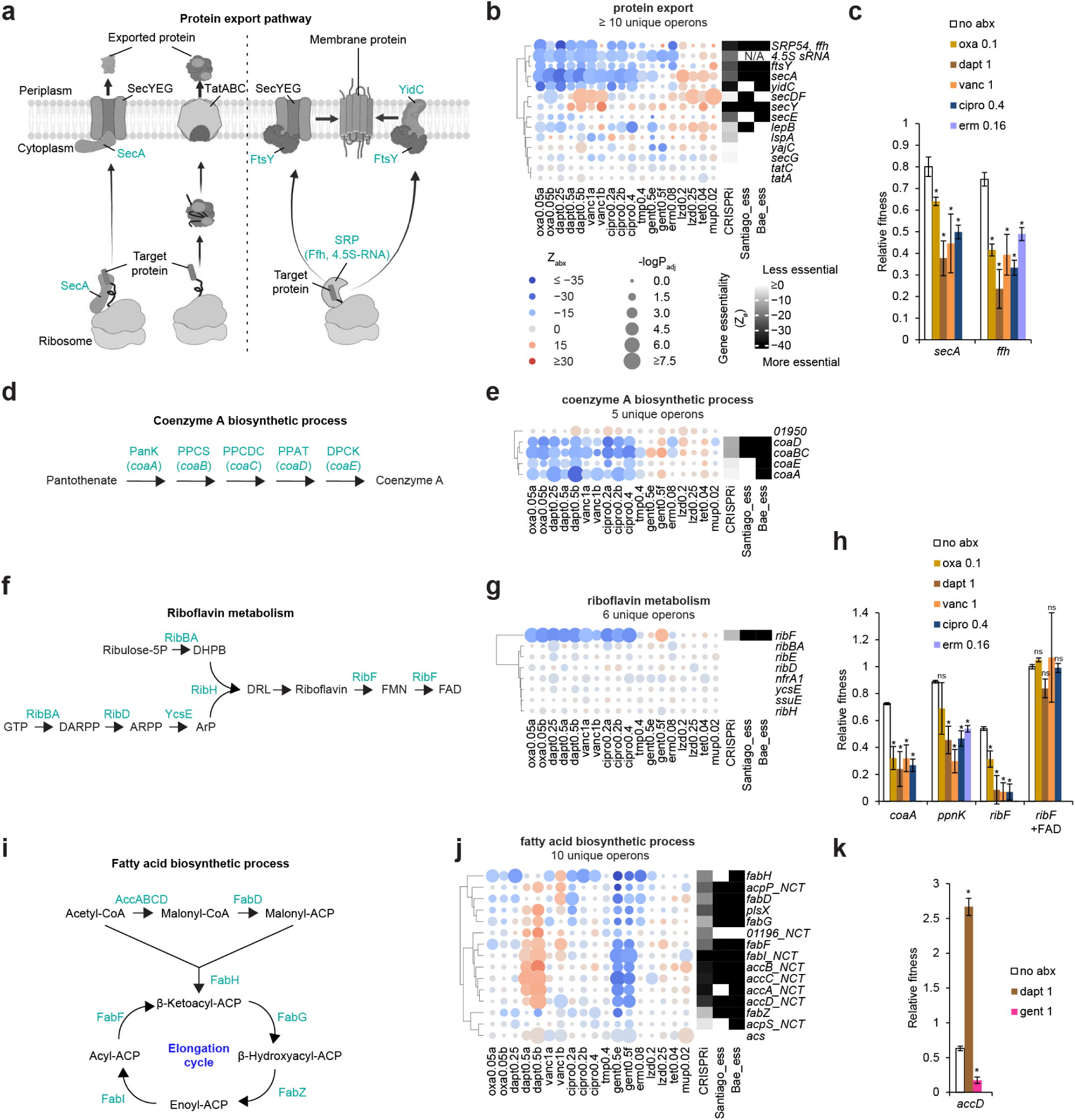
Repression of diverse biological processes sensitizes bacteria to antibiotics. (**a**) Protein export (left) and targeting to the membrane (right). Schematic is adapted from Kaushik et al^29^ with modifications. Components involved in strong sensitizing interactions are labeled in green. (**b**) Heatmap showing the relative fitness of genes in the protein export pathway (KEGG path:sao03060) in ten antibiotics as quantified by CALM-CRISPRi screens (Z_abx_) in *S. aureus* RN4220. Gene names are annotated by KEGG orthology (ko). The 4.5S sRNA was also added to the heatmap. Gene knockdowns that decreased and increased relative fitness in antibiotics are shown in blue and red circles, respectively. Size of circle indicates the negative of log-transformed adjusted P-value, -logP_adj_. Gene essentiality quantified by CRISPRi (mean of Z_Ø_ from triplicates) and qualified (binary) by Santiago’s^23^ and Bae’s^25^ Tn-seq studies are also shown. (**c**) Pairwise competition assays (Methods) measuring the relative fitness of *S. aureus* RN4220 carrying an IPTG-inducible CRISPRi system targeting *secA* and *ffh* in indicated antibiotic conditions. Concentrations of antibiotics are shown in µg/mL. Error bars indicate the standard deviation from three biological replicates. For each gene knockdown, paired t-tests were performed between the antibiotic condition and the no-antibiotic condition. * indicates P < 0.05. (**d**) Schematic of the coenzyme A biosynthesis process. (**e**) Same as (b) except heatmap is showing genes in coenzyme A biosynthetic process (GO:0015937). (**f**) Riboflavin metabolism. (**g**) Same as (b) except heatmap is showing genes in riboflavin metabolism (KEGG path:sao00740). (**h**) Same as (c) except bar plot shows the relative fitness of repression of *coaA*, *ppnK*, *ribF*, and *ribF* supplemented with 600 µM FAD. (**i**) Fatty acid biosynthetic process in bacteria. (**j**) Same as (b) except heatmap is showing genes in fatty acid biosynthetic process (GO:0006633). Genes labeled with “NCT” indicate that due to extreme gene essentiality, quantification of Z_abx_ was performed using NCT crRNAs (see “Caveat” in Methods). (**k**) Same as (c) except bar plot shows the relative fitness of repression of *accD*.

We also identified sensitizing interactions between genes and translation-targeting antibiotics, many of which were reported in previous studies. For instance, strong sensitizing interactions were observed between gentamicin and repression of ATP synthesis (arrow in **Fig. 1g** and **Fig. S4d**), as well as fatty acid synthesis (maroon label in **Fig. 1g** and **Figs. 3i, 3j**), consistent with previous findings by us and others^15,30^. Interestingly, knockdown of fatty acid synthesis led to significant increase in bacterial fitness in daptomycin (**Fig. 3j**), a new finding that was validated by our pairwise competition assay (**Fig. 3k**). Another notable hit was DNA recombination, and we confirmed that *ruvB* knockdown sensitized bacteria to multiple translation inhibitors in **Figure 2k**. We also observed sensitizing interactions involving nuclease activity (light blue label in **Fig. 1g**). Knockdown of genes encoding ribonucleases, such as *rnjA* involved in global mRNA decay^31^ and *rnr* in trans-translation^32^, has been previously shown to reduce bacterial tolerance to ribosome-targeting antibiotics in species such as *Escherichia coli* and *Bacillus subtilis*. We demonstrated that knockdown of these genes also strongly sensitized *S. aureus* to multiple translation-targeting antibiotics (genes bolded in **Fig. S4l**), including a 2- to 4-fold decrease in MICs of erythromycin, linezolid, and mupirocin (**Fig. S5x**). *rnjA* knockdown also lowered the MIC of oxacillin by 4-fold (**Fig. S5m**). Collectively, **Figure 3** and **Figure S4e – S4l** showcase diverse known and novel antibiotic-potentiating processes enriched in essential genes, establishing a genetic basis that informs the development of new combinatorial therapies (see section “Drug-drug synergy”).

### Desensitizing antibiotic-gene interactions

Repression of essential genes can not only sensitize but also desensitize bacteria to antibiotics. We found that genes encoding the transcription (eg, GO:0000428) and translation machinery (eg, GO:0006412) were prevalent among desensitizing interactions (orange and pink labels in **Fig. 1g**), especially in high concentrations of bactericidal antibiotics (dapt 0.5, vanc 1, cipro 0.4, and gent 1 in **Figs. 4a** and **4b**). Notably, even mild repression of many ribosomal genes by NCT crRNAs significantly increased relative fitness in these antibiotics (genes labeled with NCT in **Fig. 4b**), complementing the incomplete data generated from strong CT crRNAs (**Fig. S7a** and see “Caveat” in Methods). We confirmed that inhibition of *rpoC* and *rplB*, encoding subunits of RNA polymerase and ribosome, respectively, substantially increased bacterial survival in daptomycin and vancomycin (**Fig. 4c**), with *rplB* inhibition also desensitizing bacteria to ciprofloxacin and gentamicin. Repression of a subset of genes involved in energy production, including pyruvate metabolism and ETC (dot and gray labels in **Fig. 1g**, respectively), also desensitized bacteria to many bactericidal antibiotics (**Figs. S7b** and **S7c**). Together, these results showed that suppression of many essential processes, including transcription, translation, and select energy production pathways can attenuate bactericidal killing, potentially by slowing metabolism^33^ and/or inducing bacterial persistence^34-37^. In particular, our data provided strong genetic evidence supporting prior studies showing that pharmacological inhibition of transcription and translation significantly reduce antibiotic lethality^33,34^.

**Fig. 4.**
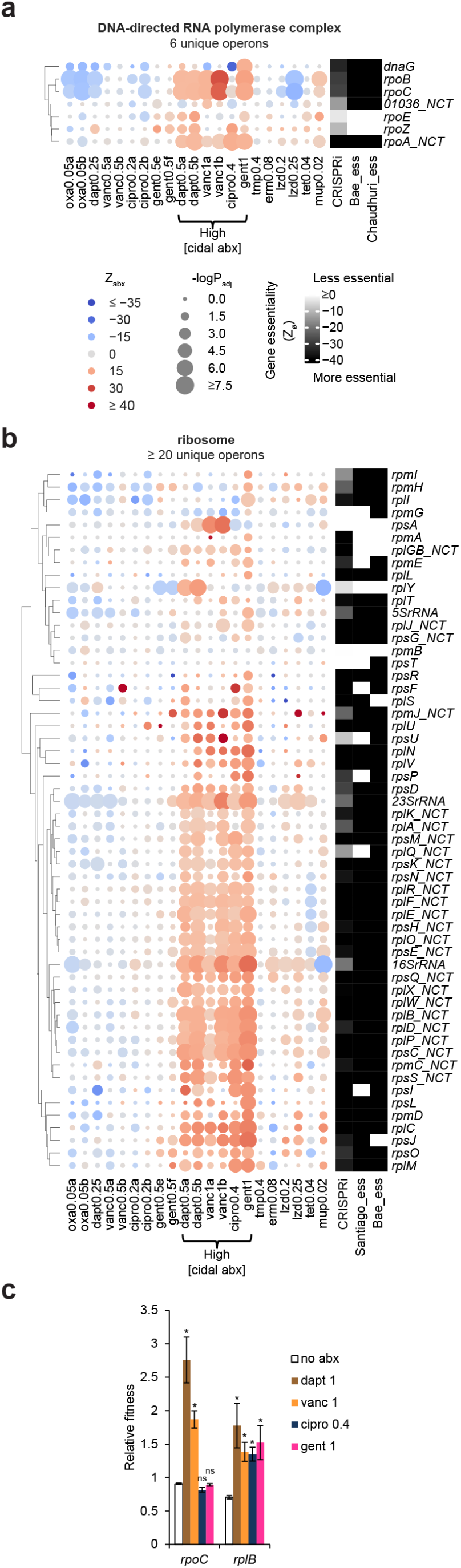
Repression of transcription and translation desensitizes bacteria to bactericidal antibiotics. (**a**) Heatmap showing the relative fitness of genes in the DNA-directed RNA polymerase complex (GO:0000428) in ten antibiotics as quantified by CALM-CRISPRi screens (Z_abx_) in *S. aureus* RN4220. The only exception is “gent1”, which shows Z_abx_ calculated from library treated with 1 µg/mL gent for 4.5 hours from a previous study^15^. Gene names are annotated by KEGG orthology (ko). Gene knockdowns that decreased and increased relative fitness in antibiotics are shown in blue and red circles, respectively. Genes labeled with “NCT” indicate that due to extreme gene essentiality, quantification of Z_abx_ was performed using NCT crRNAs (see “Caveat” in Methods). Size of circle indicates the negative of log-transformed adjusted P-value, -logP_adj_. Gene essentiality quantified by CRISPRi (mean of Z_Ø_ from triplicates) and qualified (binary) by Santiago’s^23^ and Bae’s^25^ Tn-seq studies are also shown. (**b**) Same as (a) except heatmap is showing genes in ribosome (GO:0005840). (**c**) Pairwise competition assays (Methods) measuring the relative fitness of *S. aureus* RN4220 carrying an IPTG-inducible CRISPRi system targeting *rpoC* and *rplB* in indicated antibiotic conditions. Concentrations of antibiotics are shown in µg/mL. Error bars indicate the standard deviation from three biological replicates. For each gene knockdown, paired t-tests were performed between the antibiotic condition and the no-antibiotic condition. * indicates P < 0.05.

### Gene similarity networks and genetic interactions

The genome-scale antibiotic-gene interaction profiles enabled network analysis that uncovered novel functional relationships among genes and processes. We constructed an *S. aureus* essential gene similarity network based on the interaction profiles of 440 genes and 14 antibiotic conditions (**Fig. 5a**). In a similarity network, genes displaying tightly correlated profiles form clusters, with the relative distance between clusters reflecting their functional relationships^5^. Consistent with this, we found that genes within each individual biological process had high Pearson correlation coefficients (PCCs) with each other (Methods) and were densely interconnected (**Fig. 5b**). Furthermore, several processes, including CC, DD, coenzyme A synthesis, riboflavin metabolism, and protein export, formed a discernable supercluster, which we designated Cluster I (**Fig. 5b**). In contrast, processes such as transcription (RNA polymerase), translation (ribosome), fatty acid biosynthesis, and ETC were distant from Cluster I. To corroborate this network structure, we calculated the mean PCCs among essential genes within these processes (Methods). Hierarchical clustering of inter-process PCCs of a comprehensive set of core essential processes spanning diverse functions (**Fig. 5c**) recapitulated the gene-level functional distances captured in the similarity network (**Fig. 5b**).

**Fig. 5.**
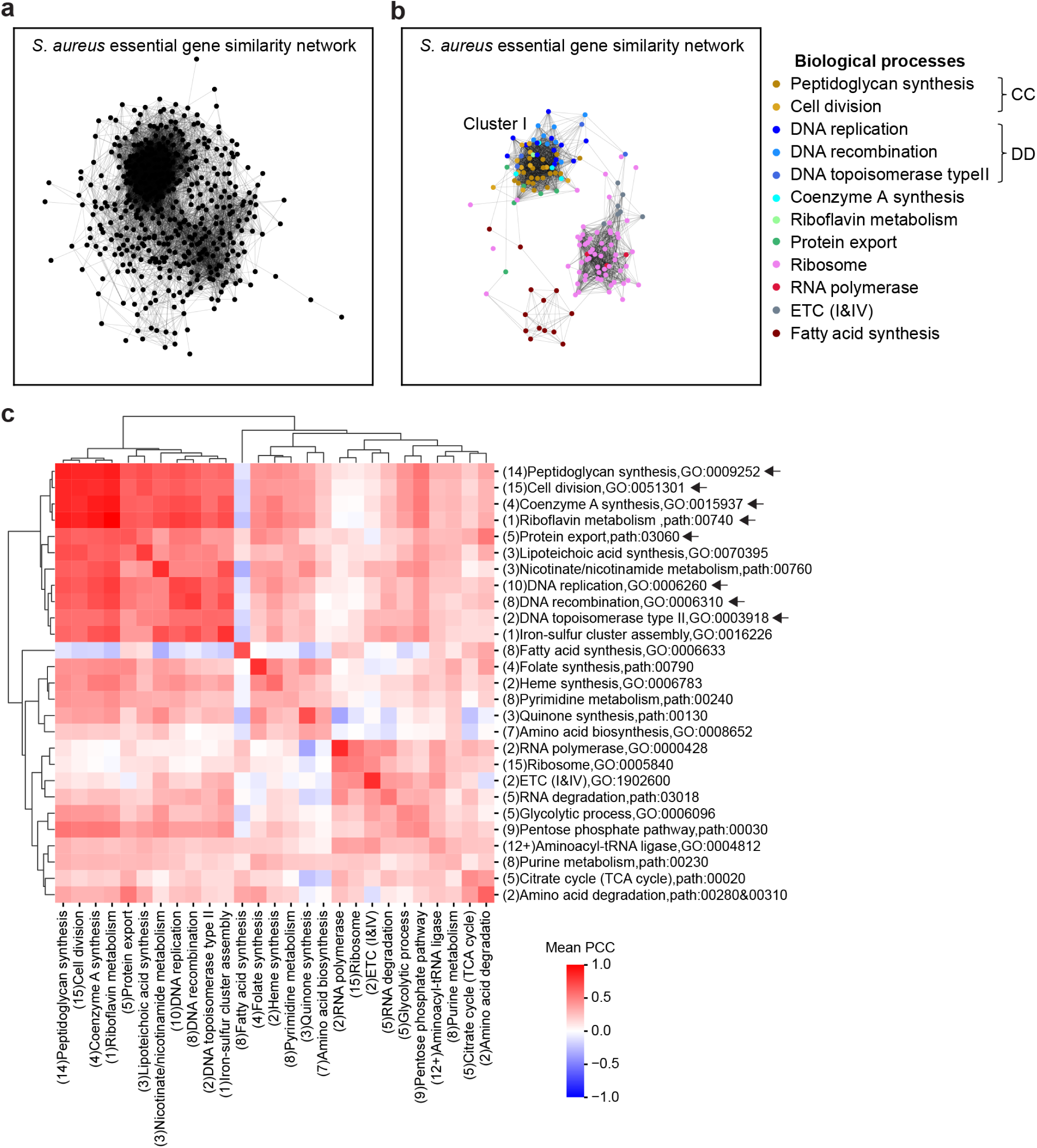
*S. aureus* essential gene similarity network. (**a**) Essential gene similarity network in *S. aureus* RN4220. Pairwise correlations between genes were calculated from the fitness profiles of 440 essential genes (defined as essential in either Santiago’s^23^ or Bae’s^25^ Tn-seq studies) in 14 antibiotic conditions. Genes with Pearson correlation coefficient (PCC) of greater than 0.65 were connected with gray edges of fixed thickness. PCC matrix and gene coordinates are given in Tables S12 and S13, respectively. (**b**) Same as (a) except only showing essential genes from select biological processes and the connections among them. (**c**) Hierarchical clustering of the pairwise mean PCCs among 27 biological processes (Methods). See also Table S14. Number of unique operons with essential genes within each process is shown in the parenthesis. Arrows indicate processes shown in Cluster I in (b).

Closely clustered genes and processes imply functional similarity. Genes with related functions tend to be enriched for synergistic genetic interactions (GIs)^5^, a type of GI in which the combined perturbation of two genes results in a significantly greater fitness defect than expected from the individual perturbations. This is because multiple partial loss-of-function mutations in the same nonlinear pathway (especially essential ones) often push the pathway below a functional threshold, aggravating fitness.

The close proximity between CC and DD genes within Cluster I (**Fig. 5b**) suggested functional similarity and potential synergies among them. To test this hypothesis, we created a dual-plasmid system enabling inducible and orthogonal transcriptional repression (**Fig. S8a**). By growing bacteria in varying concentrations of the two inducers, IPTG and xylose, we quantified fitness when pairs of CC and DD genes from ten distinct operons (**Fig. S8b**) were either individually or simultaneously repressed (**Fig. S8c**). This allowed us to calculate GI, or epistasis (ε), which measures the deviation of the observed dual perturbation effect from the expected effect based on individual perturbations (**Fig. S8d**). We observed moderate to strong synergies (ε < -0.1 and P < 0.05) in 21 of 25 tested CC-DD combinations (**Figs. 6a, 6b**), but no GIs in null controls (**Figs. S8e-S8g**). To contextualize the strength of these interactions, we referenced data from a comprehensive yeast global GI network^5^. The yeast study showed that epistasis is generally infrequent (ie, ε ≈ 0 for most GIs) and that |ε| = 0.1 represents more than one standard deviation from the mean of ∼600,000 GIs among essential genes (**Fig. S8h**). As shown in **Fig. 6a**, we found that *lexA* strongly synergized with all five tested CC genes (-0.50 < ε < - 0.15). The synergies were strong enough such that dual perturbations did not yield detectable colonies (**Fig. 6c** and *lexA* rows in **Figs. S8c** and **S8d**), even after prolonged incubation (first panel in **Fig. S8i**). Another example of strong synergy was observed between *ligA* and *murC* (**Fig. 6d**). On average, the five DD genes exhibited stronger synergies with *ftsZ* and early-stage peptidoglycan synthesis genes (*glmS*, *murC*) than with late-stage genes (*murF*, *femX*) (**Fig. 6a**). For several CC-DD pairs, the synergistic interactions were so pronounced that prolonged incubation still produced few or no colonies (arrows in **Fig. S8i**). These results strongly indicate that simultaneous genetic perturbation of diverse CC and DD genes aggravates bacterial fitness, corroborating the CC-DD synergies observed in the context of antibiotic-gene interactions (**Fig. 2**).

**Fig. 6.**
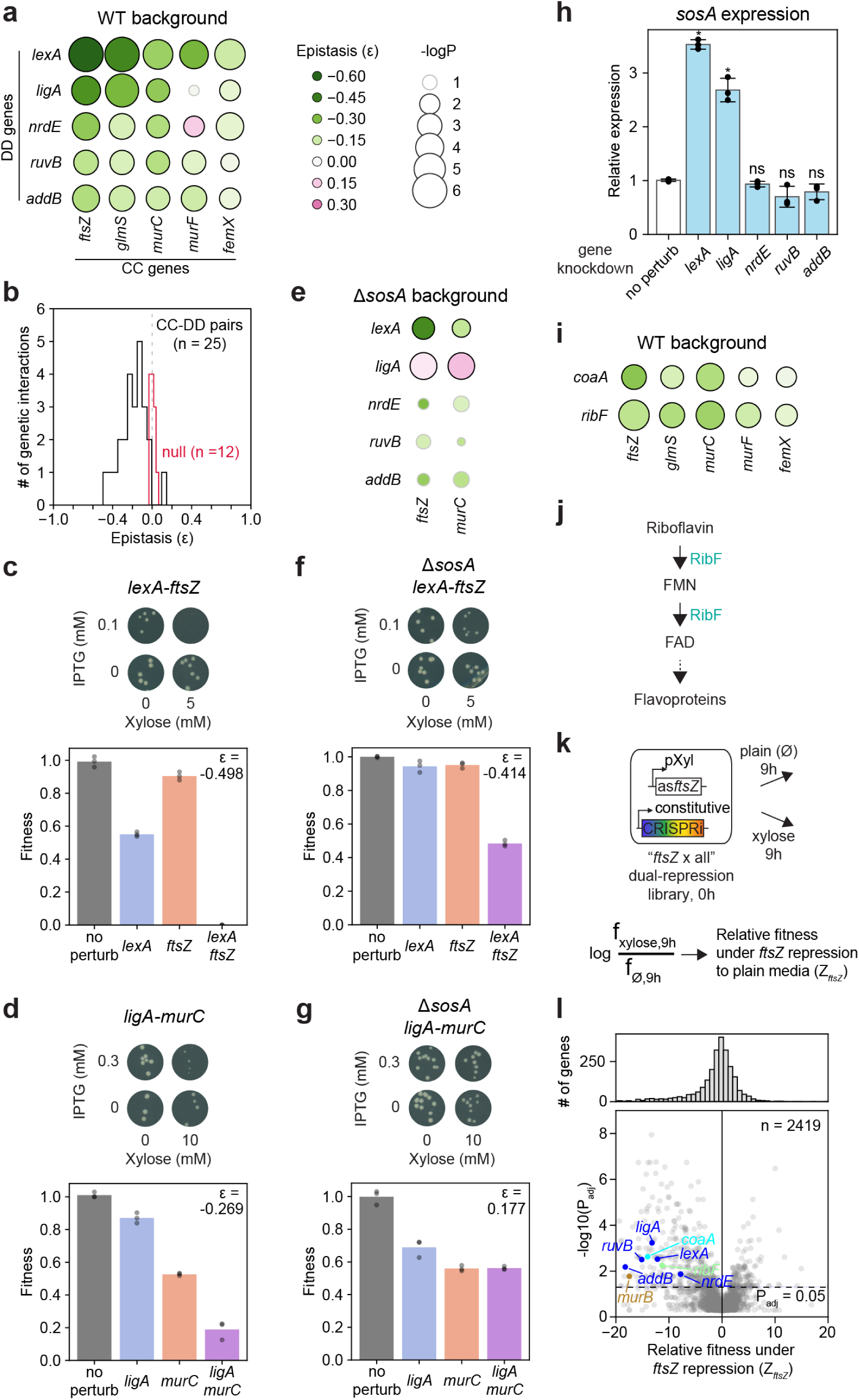
Synergistic genetic interactions. (**a**) Genetic interaction (GI) matrix between five CC and five DD genes in *S. aureus* RN4220, as quantified in Fig. S8d. For each gene-gene pair, epistasis is shown as the mean of three biological replicates. P-values were calculated using a two-sample t-test comparing the triplicates of test gene-gene pair with those of the non-targeting control induced in the same IPTG and xylose concentrations (Fig. S8f). Circles with black edges indicate P < 0.05. (**b**) Distributions of GI, or epistasis (ε) among 25 CC-DD gene pairs in (a) and baseline epistasis between plasmid backbones (null, n = 12) shown in Fig. S8f. (**c**) GI between *lexA* and *ftsZ*. Top: RN4220 cells carrying IPTG-inducible dCas9 targeting *lexA* and xylose-inducible asRNA targeting *ftsZ* were plated on TSA plates containing indicated concentrations of IPTG and xylose and grown for 22 hours at 37 °C. Bottom: Quantification of fitness (colony sizes in top panel) of bacteria that individually targeted *lexA*, *ftsZ*, or simultaneously targeted both, relative to cells with no perturbation. Epistasis between *lexA* and *ftsZ* was calculated as ε*_lexA_*_,*ftsZ*_ = W*_lexA_*, *_ftsZ_* - W*_lexA_* • W *_ftsZ_*, where W stands for fitness. ε is shown as the mean of three biological replicates. (**d**) Same as (c) except showing GI between *ligA* and *murC*. (**e**) Same as (a) except showing GI matrix between two CC and five DD genes in the Δ*sosA* background. Unlike (a), P-values were calculated using a two-sample t-test comparing the triplicates in the Δ*sosA* background with those in the wild-type background shown in (a). Circles with black edges indicate P < 0.05. See also Fig. S8j and S8k. (**f**) same as (c) except showing GI between *lexA* and *ftsZ* in Δ*sosA* background. (**g**) Same as (d) except showing GI between *ligA* and *murC* in Δ*sosA* background. (**h**) Quantification of *sosA* expression (mean ± std) by RT-qPCR. Indicated DD genes were repressed by inducible CRISPRi for 1 hour before RNA was extracted. A two-sample t-test was performed between repressed DD gene and “no perturb” control. * indicates P < 0.05. (**i**) Same as (a) except showing GI matrix between five CC genes and *coaA* and *ribF*. See also Fig. S8l and S8m. (**j**) RibF catalyzes the synthesis of FAD, an essential cofactor for flavoproteins. (**k**) Generation of “*ftsZ* x all” dual-repression library and fitness quantification. *S. aureus* cells carrying a xylose-inducible (pXyl) antisense fragment targeting *ftsZ* (as*ftsZ*) and a constitutively expressed CALM-CRISPRi library were subjected to a 9-hour outgrowth in plain or media containing xylose that mildly repressed *ftsZ*. Relative fitness of genes under *ftsZ* repression to plain media was quantified as Z*_ftsZ_*. **(l)** Volcano plot showing the relative fitness of genes under mild *ftsZ* repression to plain media, Z*_ftsZ_*. Z*_ftsZ_* values are shown in Table S15.

What is the mechanism underlying CC-DD synergies? In *S. aureus*, DNA double-strand breaks (DSBs) trigger LexA degradation, which derepresses the SOS regulon^38^, including the cell division inhibitor protein SosA^39^, a functional analog of SulA. SulA binds to FtsZ and blocks cell division^40^. We reasoned that repression of various DD processes could induce the SOS response, thereby activating SosA to inhibit cell division, which could explain the observed CC-DD synergies. To test this hypothesis, we generated a *sosA* deletion mutant and measured GIs between the five DD genes and two CC genes (*ftsZ* and *murC*). Remarkably, *sosA* deletion completely abolished the *ligA*-*ftsZ* and *ligA*-murC synergies (**Figs. 6e, 6g,** and *ligA* row in **S8k**). In contrast, CC-DD interactions involving *nrdE*, *ruvB*, and *addB* showed ε values that were not statistically different between the wild-type and Δ*sosA* backgrounds (**Figs. 6a** and **6e**). An intermediate phenotype was observed between *lexA* and CC genes. For instance, simultaneous repression of *lexA* and *ftsZ* in the Δ*sosA* background no longer prevented colony formation (**Fig. 6f**), a qualitative difference from the wild-type background (**Fig. 6c**). However, the quantified *lexA*-*ftsZ* synergy was only slightly alleviated in the Δ*sosA* background (ε = -0.498 vs. ε = -0.414, P < 0.05 in **Figs. 6c** and **6f**), suggesting the involvement of additional *sosA*-independent mechanisms. Altogether, these results indicate that *sosA* contributes to only a subset of CC-DD synergies, specifically those involving *ligA* and *lexA*. In contrast, CC-DD synergies involving *nrdE*, *ruvB*, and *addB* likely operate in a *sosA*-independent manner. Importantly, our reverse transcription quantitative PCR (RT-qPCR) experiments showed that among the five DD genes, knockdown of only *lexA* or *ligA* led to significant upregulation of *sosA* expression (**Fig. 6h**), supporting the genetic results.

Processes such as coenzyme A synthesis and riboflavin metabolism also clustered closely with CC (**Figs. 5b** and **5c**), suggesting synergistic GIs among them. This was validated by our dual-repression assays (**Figs. 6i, S8l,** and **S8m**). To explore the possible mechanism underlying the synergies between CC genes and *ribF*, we noted that RibF catalyzes the synthesis of FAD, an essential cofactor for flavoproteins (**Fig. 6j**). MurB has been validated as a flavoprotein in *S. aureus* by multiple studies^41-43^. Given MurB’s essential role in cell wall synthesis (**Fig. 2a**), its inactivation due to RibF and FAD depletion likely contributes to the observed *ribF*-CC synergy. To support this hypothesis, we generated an “*ftsZ* x all” dual-repression library and quantified the relative fitness of all genes under mild *ftsZ* repression (**Fig. 6k**). This global assay showed that among all flavoprotein-encoding genes, *murB* repression significantly reduced relative fitness when *ftsZ* was simultaneously repressed (**Figs. S9a** and **S9b**). Moreover, the “*ftsZ* x all” screen also showed that repression of *coaA*, *ribF*, and the five DD genes significantly reduced the relative fitness under mild *ftsZ* repression (**Fig. 6l**), further supporting their synergies with *ftsZ*. Collectively, our CRISPRi screens and dual-repression assays uncovered new synergistic GIs between CC and coenzyme A synthesis, as well as CC and riboflavin metabolism, suggesting functional links between these processes.

### Drug-drug synergies

To investigate if we can translate synergistic GIs into viable combinatorial drug therapies, we identified SMIs targeting products of genes in Cluster I, including RuvAB^44^ (DNA recombination), SecA^45^ (protein export), coenzyme A biosynthesis^46^, and RibF^47^ (riboflavin metabolism), and tested if they could synergize with the antibiotics used in this study. Quantifying antibiotic-SMI interactions by solely measuring the changes in MIC (ΔMIC) at a terminal growth time point (∼24 hour) could be biased. To be more stringent, we also calculated epistasis (ε) between antibiotics and SMIs over an extended period of growth (Methods), and only considered a simultaneous reduction in MIC (ΔMIC ≤ ½) and strong negative epistasis (ε ≤ -0.4) to be synergistic antibiotic-SMI interactions. Of the 20 antibiotic-SMI combinations tested on *S. aureus* RN4220, 11 exhibited synergy (**Figs. 7a** and **S10a-o**). Critically, four SMIs also strongly synergized with antibiotics in *S. aureus* USA300 (**Figs. 7b-7e** and **S10p-v**), a MRSA strain^22^ with multiple resistance determinants, including methicillin, ciprofloxacin, erythromycin, tetracycline, and mupirocin. Daptomycin, an antibiotic commonly used for treating MRSA infection, strongly synergized with three SMIs targeting distinct processes (**Fig. 7b**), including methylbenzethonium chloride (MTBC) that targets RibF (**Fig. 7c**) and visomitin that targets RuvAB (**Fig. 7d**). Among all antibiotic-SMI interactions, the strongest synergy was observed between gentamicin and 5-(Tetradecyloxy)-2-furoic acid (TOFA) (**Fig. 7e**), which targets the AccABCD complex^48^ in the fatty acid synthesis pathway (**Fig. 3i**). This finding is consistent with our antibiotic-gene interaction result (**Fig. 3j, 3k**) and our previous work^15^. These results demonstrate the power of CRISPRi functional genomics in identifying new antibiotic-synergizing essential gene targets and the potential to translate them into effective drug combinations. However, we caution that drug-drug synergy can also arise from drug pleiotropy or from changes in cellular permeability that alter intracellular drug concentrations without affecting their targets^49,50^. Consequently, observed drug-drug interactions may not always reflect GIs identified through genetics and functional genomics studies.

**Fig. 7.**
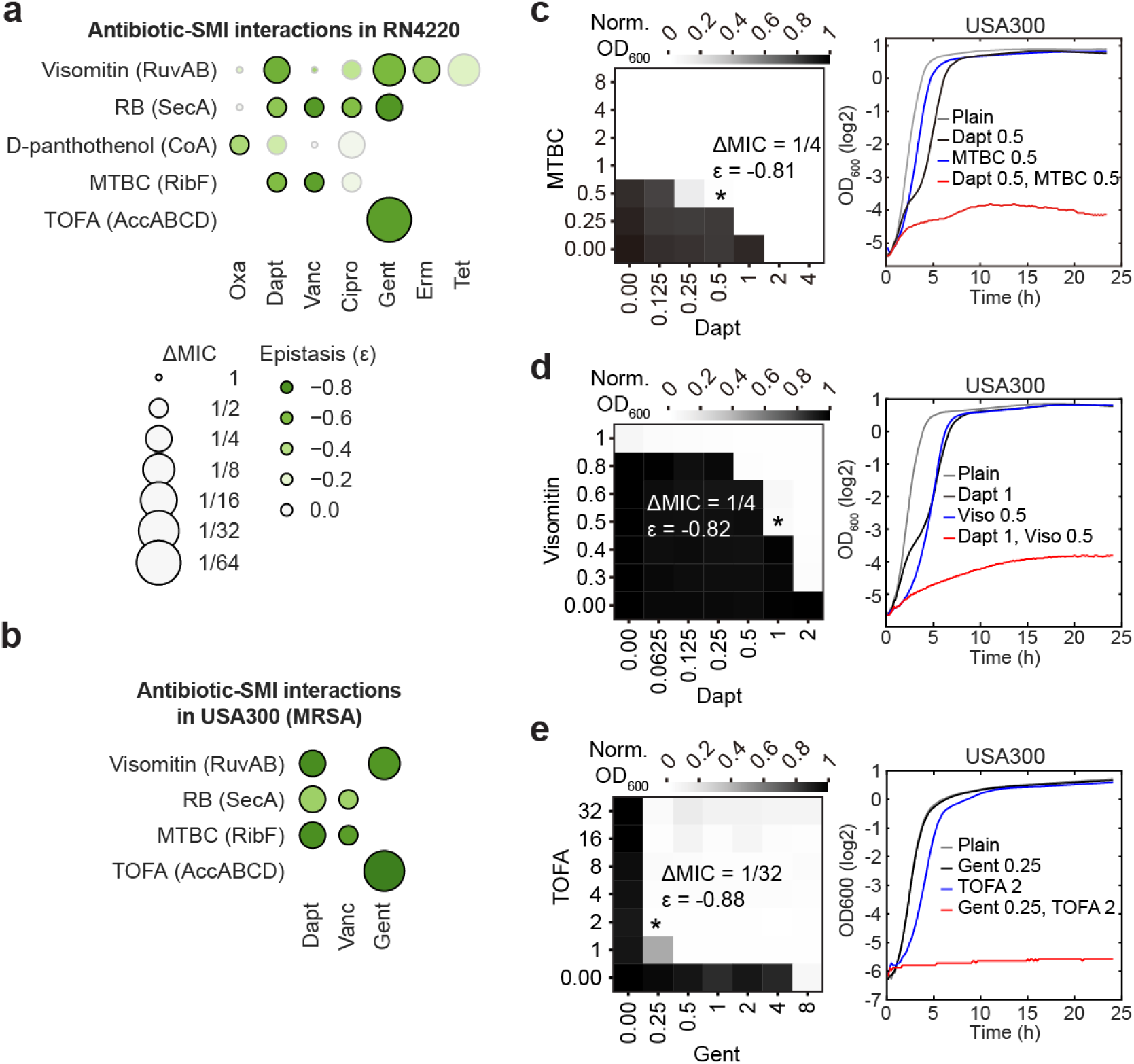
Drug-drug synergy. (**a**) Interactions between antibiotics (x-axis) and SMIs (y-axis) in *S. aureus* RN4220 quantified by checkerboard assays. Targets of SMIs are shown in parentheses. Changes in MIC (ΔMIC) and epistasis (ε) were quantified. Synergistic antibiotic-SMI combinations (ΔMIC ≤ ½ and ε ≤ -0.4) have black edges while other combinations have gray edges. Abbreviations are as follows. RB: rose bengal. MTBC: methylbenzethonium chloride. TOFA: 5-(Tetradecyloxy)-2-furoic acid. (**b**) Same as (a) except showing antibiotic-SMI interactions in *S. aureus* USA300. (**c**) Left: checkerboard assay of daptomycin-MTBC combination for *S. aureus* USA300, showing bacterial growth at various concentrations of antibiotic and SMI, normalized to bacterial growth in plain media (ie, 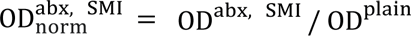) 22-24 hours post inoculation. An asterisk “*” labels the well where the change in MIC (ΔMIC) and epistasis (ε) were calculated (Methods). Concentrations of SMI and antibiotic are in μg/mL. Right: four growth curves related to the well labeled with asterisk: plain media (gray), antibiotic alone (black), SMI alone (blue), and both antibiotic and SMI combined (red). (**d, e**) Same as (c) except showing the daptomycin-visomitin and gentamicin-TOFA combinations for *S. aureus* USA300, respectively.

## DISCUSSION

Motivated by the scarcity of systems-level studies of gene vulnerability under antibiotic stress in *S. aureus*, we quantified global gene fitness across ten antibiotics with various modes of action by leveraging our previously developed CALM-CRISPRi functional genomics platform. Our approach revealed a comprehensive set of known and novel biological processes affecting bacterial fitness under antibiotic stress. Importantly, essential genes dominated the hundreds of significant antibiotic-gene interactions we identified, a finding not manifested by previous transposon-based studies. Recent CRISPRi work in *M. tuberculosis* and GI study in yeast also found that essential genes participated in significantly more antibiotic-gene^6^ and gene-gene interactions^5^ than nonessential genes, supporting our findings.

Inhibition of diverse essential processes could both sensitize and desensitize bacteria to antibiotics. The essential CC, DD, coenzyme A biosynthesis, protein export, and riboflavin metabolism pathways were among the most vulnerable biological processes under many antibiotic conditions, especially those that target cell wall/membrane and DNA synthesis. Among these processes, we found genes, such as *ruvB* and *rnjA*, sensitized *S. aureus* to additional antibiotics that target translation (**Figs. 2g, 2k, S4l,** and **S5x**), making these gene products new attractive targets for combinatorial antimicrobial therapy. Perturbation of essential genes could also desensitize bacteria to antibiotics. We found that CRISPRi repression of genes involved in transcription, translation, and selective energy processes tended to antagonize bactericidal killing by many antibiotics, such as daptomycin, vancomycin, ciprofloxacin, and gentamicin. These findings establish the genetic basis for previous observations showing that pharmacological inhibition of transcription, translation, and proton motive force slowed cellular metabolism and reduced antibiotic lethality^33,34^.

The genome-scale antibiotic-gene interaction profiles led us to construct an essential gene similarity network (**Figs. 5a** and **5b**), which revealed both known and novel functional relationships among biological processes. We observed extensive synergistic interactions between diverse CC and DD processes (**Figs. 6a** and **6b**). While previous mechanistic studies^40,51^ focused on characterizing the connections between cell division (eg, *sulA* and *ftsZ*) and DNA replication (eg, *dnaA* and *dnaB*), the scale of our approach identified a more extensive CC-DD network encompassing genes in various stages of cell wall synthesis (eg, *glmS* and *murC*), nucleotide synthesis (eg, *nrdE*), and DNA recombination (eg, *addB* and *ruvB*). Critically, our dual-perturbation assay provided direct evidence that simultaneous transcriptional repression of these essential CC and DD genes aggravated bacterial fitness, which has not been shown before. Our genetic analysis further demonstrated that CC-DD synergies could be either dependent on, or independent of SosA, a functional analog of the cell division inhibitor SulA in *S. aureus*. SosA contributed significantly to CC-DD synergies involving *lexA* or *ligA*, which are genes whose perturbation also strongly induced SosA. In contrast, deletion of *sosA* did not significantly reduce CC-DD synergies involving the other three tested DD genes: *nrdE*, *ruvB*, and *addB*, suggesting that other pathways may underlie these interactions. An alternative explanation is that chromosome replication and cell division are two highly coordinated and essential processes in bacteria^51-53^, and multiple partial loss-of-function mutations within this coordination may simply push the system below a functional threshold and aggravate fitness without necessarily involving additional pathways. An essential gene similarity network constructed in *B. subtilis* based on chemical-genomic screens (35 compounds x 289 genes) also revealed close proximity between CC and DD processes^54^, supporting our results in *S. aureus*. Interestingly, unlike these findings in bacteria, the comprehensive yeast GI study did not report prominent CC-DD synergies^5^. This seems to be congruous with a fundamental distinction in eukaryotes, where chromosome replication (S phase) and cell division (M phase) are temporally separated. In addition to the extensive CC-DD synergies, we also identified and validated new synergistic GIs involving CC, coenzyme A synthesis, and riboflavin metabolism, suggesting functional links among them.

The findings from our antibiotic-gene and gene-gene interaction assays establish a genetic basis that informs the development of new combinatorial therapies against *S. aureus*. Notably, 11 out of 20 antibiotic-SMI combinations we tested exhibited potent killing (**Figs. 7a** and **7b**). In particular, novel antibiotic-SMI combinations involving visomitin, MTBC, and TOFA were highly synergistic when tested on MRSA USA300. Visomitin and MTBC target RuvAB and RibF, respectively, which are molecular targets that do not have approved antibiotics. Another potentially attractive drug target with no known inhibitors is Ribonuclease J1, encoded by *rnjA*. Our results, along with a previous study^31^, show that *rnjA* repression sensitized *S. aureus* and *B. subtilis* to antibiotics of various modes of action, including those targeting the cell wall/membrane, DNA synthesis, and translation. Together, these results provided a strong rationale for developing antimicrobial drugs against these gene targets, and for deploying them in combination therapies as a viable antimicrobial strategy.

Our study represents the most comprehensive investigations of the genetic landscape of antibiotic sensitivity in *S. aureus* to date. The highly sensitive CALM-CRISPRi platform enabled fitness profiling of essential genes under antibiotic conditions that were not possible with Tn-seq. Our small-scale similarity network analysis based on 440 essential genes and 14 antibiotics identified novel GIs within the core essential bacterial genome, new drug targets, and rational drug synergies. These findings encourage a more comprehensive investigation of GIs across the entire genome, which could significantly advance our understanding of bacterial genetic architecture and cellular organization, and accelerate antimicrobial discovery. Excitingly, two recent studies employed CRISPRi genomics to investigate GIs in *Streptococcus pneumoniae*^55^, encompassing ∼50% of possible digenic combinations, and in *B. subtlis*^56^, focusing on digenic combinations involving genes in CC. We hope that our work, along with these GI studies, will motivate future efforts to explore novel biological principles in diverse bacterial species through high-throughput CRISPR genomics.

## ACKNOWLEDGEMENTS

We thank the Jiang lab and Tavazoie lab for insightful comments on the project. We thank Christopher D. Johnston for kindly providing the xylose-inducible plasmid, pEPSA5. This work was supported by NIH grants R00AI153530 (WJ), R01AI077562 (ST), R01GM139215 (ST) and seed fund from Icahn School of Medicine at Mount Sinai.

## AUTHOR CONTRIBUTIONS

WJ conceived the study. WJ and WL performed wet-lab experiments and data analysis. SB, FP, JH, JG, and QH assisted with cloning, DNA extraction, PCR and growth measurements. WJ performed dry-lab analysis, with assistance from ML, PO and HC. WJ and ST secured funding for the study. WJ wrote the paper.

## DATA AVAILABILITY

Raw sequencing data are deposited to NCBI SRA under project number PRJNA1181893. CRISPRi sequencing results and analyses are available in Tables S2-S15.

## MATERIALS AND METHODS

### Bacterial strains and reagents

Cultivation of *Staphylococcus aureus* strains RN4220^20^, USA300^57^, JE2^21^ were carried out in tryptic soy broth (TSB) medium (BD) at 37 °C with shaking (rpm 220), or on tryptic soy agar (TSA) plates. Whenever necessary, medium was supplemented with appropriate antibiotics to select for plasmid transformation. In these cases, the concentrations of antibiotics were as follows: chloramphenicol, 5 µg/mL; erythromycin, 10 µg/mL; tetracycline, 5 µg/mL. Antibiotics, SMIs, and key reagents used in the study are shown in **Table S16**.

### Plasmid cloning

Plasmid pWJ584 was constructed by Gibson assembly of two PCR products. One PCR was performed using plasmid pWJ402 as template and primers W1956 and W1948. Another PCR was performed using pWJ420 as template and primers W1957 and W1961.

Plasmid pWJ587 (chloramphenicol-resistant) was constructed by Gibson assembly of two PCR products. One PCR was performed using plasmid pWJ418 as template and primers W1341 and W1342. Another PCR was performed using pWJ444 as template and primers W1971 and W1972. The canonical CRISPR array has the structure “repeat-spacer-repeat”. pWJ587 contained two BsaI cleavage sites between the two repeats, facilitating spacer cloning. Spacer is the DNA precursor to crRNA.

Plasmid pWJ676 was constructed by Gibson assembly of three PCR products. The first PCR was performed using pT181 as template and primers W2326 and W666. The second PCR was performed using pWJ584 as template and primers W2327 and W2328. The third PCR was performed using pWJ584 as template and primers W2329 and W2331.

Similar to previous studies^14,15^, CRISPR spacer cloning was performed by ligation of annealed oligonucleotide pairs and BsaI-digested parent vector, pWJ444 (constitutive dCas9) or pWJ587 (IPTG-inducible dCas9). Plasmids pWJ402, pWJ418, pWJ420, pWJ444 were previously published^15^.

pWJ700 harbored a single repeat as opposed to “repeat-spacer-repeat”, but was otherwise isogenic to pWJ587. To obtain pWJ700, we streaked RN4220 cells harboring an IPTG-inducible *ftsZ*-targeting CRISPRi plasmid, pWJ628, on TSA containing 10 µg/mL chloramphenicol and 1 mM IPTG. As *ftsZ* is an essential gene, it was fairly easy to isolate escaper cells containing mutated plasmid that had lost *ftsZ*-targeting spacer through homologous recombination of the flanking repeats. Sequence of pWJ700 was confirmed by sequencing.

To perform genetic complementation of *ruvB* and *ligA*, it was necessary to express them containing synonymous mutations that negate CRISPRi targeting on a second plasmid. Plasmid pWL15, a *ruvB* complementation plasmid, was constructed via Gibson assembly of three PCR products. The first PCR product was generated using plasmid pWJ418 as template with primers W2286 and W2287. The second PCR product was generated using RN4220 genomic DNA as template with primers WL1 and WL4. The third PCR product was generated using RN4220 genomic DNA as template with primers WL2 and WL3. Similarly, plasmid pWL25, a *ligA* complementation plasmid, was constructed through Gibson assembly of three PCR products. The first PCR product was generated using plasmid pWJ418 as template with primers W2286 and W2287. The second PCR product was generated using RN4220 genomic DNA as template with primers WL5 and WL8. The third PCR product was generated using RN4220 genomic DNA as template with primers WL6 and WL7.

To perform genetic interaction assays, it was necessary to change the resistance of pWJ587-based plasmids targeting various genes from chloramphenicol to tetracycline. This was achieved by Gibson assembly of two PCR products. One PCR was performed using plasmid pT181 as template and primers W1311 and W1312. Another PCR was performed using the pWJ587-based plasmid as template and primers W2394 and W2395. For example, the chloramphenicol-resistant cassette of pWJ700 (single-repeat) was changed to tetracycline in this manner, generating pWL17.

Antisense RNA fragments targeting genes of interest were cloned after a xylose-inducible promoter in a pC194-based plasmid, pEPSA5^58^. To do so, pEPSA5 was linearized by EcoRI. Next, PCR was performed to amplify an antisense fragment targeting genes of interest in RN4220. The two fragments were ligated by Gibson assembly. Sequences of antisense fragments and primers used for cloning were listed in **Table S17**. All PCRs and Gibson assembly reactions were performed using Q5^®^ High-Fidelity DNA Polymerase and NEBuilder^®^ HiFi DNA Assembly Master Mix supplied by NEB, respectively.

### Construction of the Δ*sosA* mutant strain

To delete the *sosA* (SAOUHSC_01334) in *S. aureus* RN4220, we took an allelic exchange approach facilitated by CRISPR counterselection. First, approximately 1-kb regions upstream and downstream of *sosA* were amplified by PCR using primer pairs WL97/WL98 and WL99/WL100, respectively. The plasmid backbone of vector pWJ244 was amplified using primers W1005 and W1055. The three fragments were assembled via Gibson assembly to generate the allelic exchange plasmid pWL85, which was then transformed into *E. coli* DH5α for propagation and subsequently purified. Next, pWL85 was introduced into *S. aureus* RN4220 and transformants were selected on TSA containing chloramphenicol (10 µg/mL) at 37 °C. As pWL85 did not contain *S. aureus* origin of replication, chloramphenicol-resistant clones were exclusively integrants. Next, to promote plasmid excision, we infected integrants with a temperature-sensitive, erythromycin-resistant phagemid pWJ326, which carries a CRISPR-Cas9 system targeting the chloramphenicol-resistant cassette on pWL85. Infected cells were plated on TSA containing erythromycin (5 µg/mL) and incubated at 30 °C to select for plasmid excision. Finally, successful deletion of *sosA* was confirmed by PCR using primer pairs WL101/W1043 and W614/WL102, and further validated by Sanger sequencing. The resulting RN4220ΔsosA strain was designated LIVA6.

### Generation of genome-scale CRISPRi libraries using CALM and subsequent selections in antibiotic conditions or xylose

Method for generating genome-scale CRISPRi libraries in *S. aureus* is similar to our previous study^15^. The CALM-CRISPRi machinery is encoded on a single plasmid, pWJ584, which contains *tracr*, hd*cas9*, an empty CRISPR array (denoted as “R”), *cas1*, *cas2*, *csn2* (**Fig. 1a**) and a chloramphenicol-resistant cassette. The CRISPR adaptation machinery (*cas1*, *cas2* and *csn2*) is under an IPTG-inducible promoter, pSpac). CRISPR components necessary for transcriptional repression (ie, hdCas9, crRNA and tracrRNA) are constitutively expressed. To start, a single colony of *S. aureus* RN4220 or JE2 strains harboring pWJ584 was grown overnight in 4 mL of TSB with chloramphenicol (5 µg/mL). Culture was diluted 1:200 in 15 mL of fresh TSB (no antibiotics) with 2 mM IPTG to induce the expression of Cas1, Cas2 and Csn2 and grown until OD_600_ reached 1.0 (typically 3 – 4 h). To make competent cells, cells were pelleted and washed two times using one volume of sterile water at room temperature. Cells were ultimately re-suspended in 1/100^th^ volume of sterile water. It is crucial to perform these steps at room temperature and avoid cold shocking bacterial cells, as we observed that cells prepared at 4 °C contained libraries with considerably more spacers matching the plasmid rather than the chromosome.

50 µL of competent cells were mixed with 2 µg of sheared genomic DNA (to an average of 150 bp) prepared from the same bacterial host and incubated 5 min at room temperature. Electroporation was performed using MicroPulser (Bio-Rad) with the default staph program (2 mm, 1.8 kV and 2.5 ms). Of note, we found ∼50% of the cells were killed by electroporation, which was moderate. After electroporation, cells were immediately re-suspended in 500 µL of TSB and recovered at 37 °C for 15 min with shaking. Next, 200 µL of recovered cells were transferred to 15 mL of pre-warmed TSB with chloramphenicol (5 µg/mL) and recovered for an additional 5 h at 37 °C with shaking. During this recovery period, CRISPR adaptation occurred, and the resulting culture, typically at 3 – 4 in OD_600_ (∼3.0 or 10^9^ CFU/mL, constituted the genome-scale CRISPRi library we deemed as Time 0 h in **Fig. 1a**. Typically, 7 mL of the library culture was pelleted, lysed, amplified by PCR. Products were purified by AMPure XP beads or Axygen™ AxyPrep MAG PCR Clean-Up Kit and sent for Illumina sequencing. We found the two types of beads to be interchangeable.

To profile the genetic fitness landscape of *S. aureus* in antibiotics, 7 mL of CALM-CRISPRi libraries were transferred to a 2 L Erlenmeyer flask containing 700 mL of plain TSB (pre-warmed at 37 °C), or TSB containing sublethal concentrations of tested antibiotics, including oxacillin (0.05 µg/mL), daptomycin (0.25 and 0.5 µg/mL), vancomycin (1 µg/mL), ciprofloxacin (0.2 and 0.4 µg/mL), gentamicin (0.5 µg/mL), trimethoprim (0.4 µg/mL), erythromycin (0.08 µg/mL), tetracycline (0.04 µg/mL), linezolid (0.2 and 0.25 µg/mL), and mupirocin (0.02 µg/mL). Chloramphenicol was not added during this selection to avoid potential drug-drug interactions. Whenever daptomycin was used, media was also supplemented with 1 mM CaCl_2_. Cultures were incubated at 37 °C with shaking (140 rpm) for 9 hours, reaching an OD_600_ of 4-5. Finally, 7 mL of the culture was pelleted, lysed, amplified by PCR and sent for Illumina sequencing as described in section “*S. aureus* Harboring CRISPR Adaptation Machinery (hdcas9) Generates Single-spacer (1S) Libraries Targeting *S. aureus*” in a previous publication^15^.

To generate the “*ftsZ* x all” dual-repression libraries, *S. aureus* RN4220 cells harboring pWJ633 (encoding an inducible antisense fragment targeting *ftsZ*) and pWJ676 (CALM) were used, following the same procedure as described for *S. aureus* carrying pWJ584. pWJ676 is identical to pWJ584, except that the chloramphenicol-resistance cassette is replaced with a tetracycline-resistance cassette. After constructing the “*ftsZ* x all” dual-repression library, 7 mL of the library culture was transferred to a 2 L Erlenmeyer flask containing 700 mL of TSB containing 5 µg/mL of chloramphenicol (pre-warmed at 37 °C), or TSB containing 5 µg/mL of chloramphenicol plus 10 mM xylose. Chloramphenicol was used to maintain pWJ633, and xylose was used to induce antisense-*ftsZ* expression. Cultures were incubated at 37 °C with shaking (140 rpm) for 9 hours, reaching an OD_600_ of 4-5, equivalent to ∼7 generations. Finally, 7 mL of the culture was pelleted, lysed, amplified by PCR and sent for Illumina sequencing as described in section “*S. aureus* Harboring CRISPR Adaptation Machinery (hdcas9) Generates Single-spacer (1S) Libraries Targeting *S. aureus*” in a previous publication^15^.

### Sequencing analysis and gene fitness quantification

Illumina sequencing data was processed as described in section “Spacer identification and sequence alignment” under “Data Analysis of Single-spacer (1S) and ‘‘One-vs-all’’ Libraries” of a previous publication^15^. Briefly, the sequences of spacers, which are the DNA precursors to crRNAs, were identified and aligned to the genomes of *S. aureus* NCTC8325 (NCBI reference genome: NC_007795.1) and USA300 (NCBI reference genome: NC_007793.1) using Burrows-Wheeler Alignment tool^59^. A key difference between RN4220 and its parental strain, NCTC8325, is the absence of three prophages (SAOUHSC_01514 – SAOUHSC_01582, SAOUHSC_02016 – SAOUHSC_02089, and SAOUHSC_02164 – SAOUHSC_02239). For each sample, the number of reads for each spacer was recorded. Throughout this study, the terms ‘‘crRNA’’ and ‘‘spacer’’ are used interchangeably.

Our work quantified both gene fitness in plain TSB (Z_Ø_) and their relative fitness in antibiotics (Z_abx_). Quantification of gene fitness in plain TSB (Z_Ø_) is described as follows. For each CT crRNA, the normalized frequency of it, 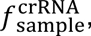 was calculated by dividing its reads by the total number of reads in that sample. The fitness of each crRNA in plain TSB media (Ø) was calculated as the log2-transformed fold-change of frequency of crRNA measured at 9 hour to that measured at 0 hour:

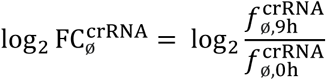

crRNA positions with reads below 10 in both the numerator (ie, 9-hour) and the denominator (ie, 0-hour) were excluded from analysis. For each gene, we quantified 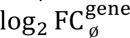 by averaging all remaining CT crRNAs targeting that gene. To minimize potential off-target effects, we ranked all 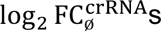 in ascending orders (ie, from 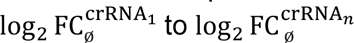) and removed the CT crRNAs with the lowest and the highest values, unless the gene has three or fewer CT crRNAs. This gives:

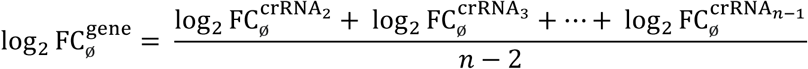

Subsequently, 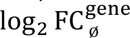 was standardized by Z-scoring, using pseudogenes made up by null crRNAs. Our CALM-CRISPRi platform routinely generated highly comprehensive crRNA libraries covering ∼95% of all targetable sites in the genome, encompassing diverse genetic elements and intergenic regions. We considered sequences not annotated by NCBI reference genomes but are 100 bp upstream from operon start codons as intergenic regions. Approximately 4,500 crRNAs targeting these intergenic regions were assigned as null crRNAs. For each treatment condition, fifty random null crRNAs were chosen to create a pseudogene, and fitness of the pseudogene, 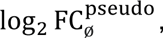 was calculated by averaging these null crRNAs. Next, a collection of fifty pseudogenes were created in this manner, and the mean and standard deviation of fifty 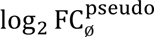 s were calculated as 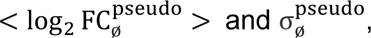 respectively. This allowed us to calculate Z-scores for each gene:

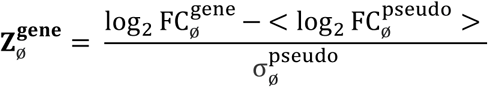

This standardized Z-score, serving as the final gene fitness score, allows samples to be compared across all tested conditions, including plain TSB media (Ø) and antibiotic treatments (abx). For each gene, a Mann-Whitney U test was performed between all CT crRNAs targeting the gene and the null crRNAs of the pseudogenes. The resulting P-value is adjusted by the Benjamini-Hochberg method.

Similarly, the relative gene fitness in antibiotics compared to plain media can be also quantified as follows:

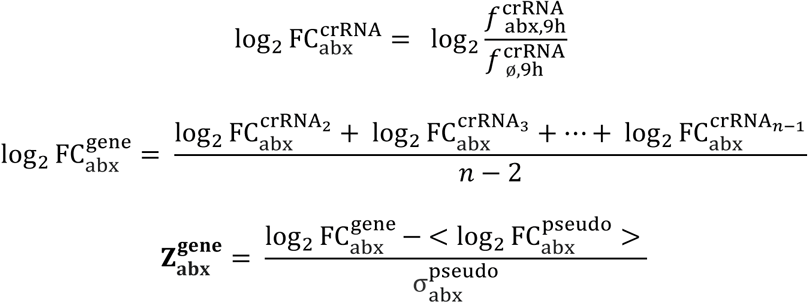

Similarly, crRNA positions with reads below 10 in both the numerator (ie, abx, 9-hour) and the denominator (ie, Ø, 9-hour) were excluded from analysis. Throughout, 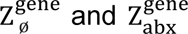 have been simplified as Z_Ø_ and Z_abx_, respectively.

In this study, we generated at least seven independent CALM-CRISPRi libraries in *S. aureus* RN4220. Libraries C2821, C2924, and C2998 (**Fig. S1e**) were chosen as the three biological replicates to calculate mean Z_Ø_ values.

### Caveat

We also quantified gene relative fitness in antibiotics using the mildly-targeting NCT crRNAs (**Table S4**). For many CT crRNAs targeting highly essential genes, such as those involved in ribosome, RNA polymerase, and fatty acid synthesis, their raw sequencing counts at 9-hour dropped to a very small number (<10) in both plain media and antibiotic conditions (**Fig. 1a**), preventing reliable quantification of Z_abx_ (eg, P_adj_ = 1 for many ribosomal genes in **Fig. S7a**). In contrast, the mild NCT crRNAs did not severely deplete at 9-hour, offering a better dynamic range for quantifying the relative fitness of these highly essential genes in antibiotics. We used NCT crRNAs to quantify a portion of highly essential genes encoding the ribosome (**Fig. 4b**), RNA polymerase (*rpoA* and *SAOUHSC_01036* in **Fig. 4a**), and fatty acid synthesis (*acpPS*, *accABCD*, *fabI*, and *SAOUHSC_01196* in **Fig. 3j**).

The general flow of quantification is the same as before except that NCT, instead of CT crRNAs were used for each gene:

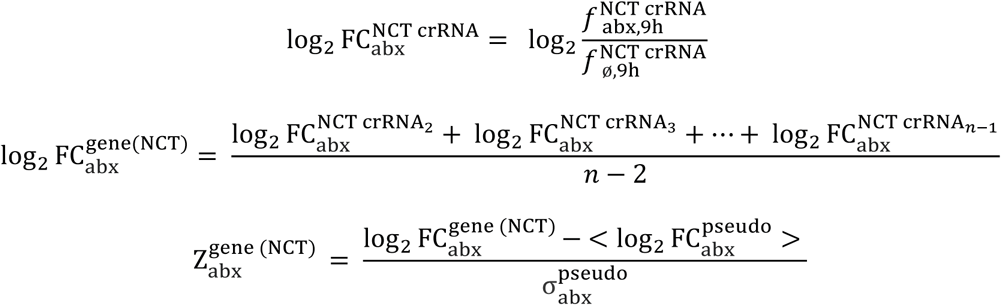

### Functional enrichment analysis

Functional enrichment analysis was performed using iPAGE, an information-theoretic pathway analysis tool^60^ we previously developed. We used both KEGG pathway (https://www.genome.jp/kegg-bin/show_organism?menu_type=pathway_maps&org=sao) and GO term annotations (https://biocyc.org/GCF_000013425/organism-summary) for *S. aureus* NCTC 8325, a parental strain for RN4220. Overall, ∼37% and ∼83% of the RN4220 genome is annotated by KEGG pathways and GO terms, respectively. While *S. aureus* genes are poorly annotated by KEGG pathways (eg, well known genes such as *ftsZ*, *lexA*, and *addAB* have no annotated pathways), GO terms tend to over-annotate, which can also lead to dilution of significant pathways. Discrete mode of iPAGE was used for enrichment analysis. Genes with Z_abx_ ≤ 9 and P_adj_ < 0.05 were categorized as strongly decreased relative fitness in antibiotics while those with Z_abx_ ≥ 9 and P_adj_ < 0.05 were categorized as strongly increased relative fitness in antibiotics. For antibiotic conditions that have biological replicates, mean Z_abx_ values were used for enrichment analysis.

### Pairwise competition assay

*S. aureus* RN4220 cells harboring an IPTG-inducible dCas9, along with no spacer (pWJ700) and spacers targeting genes of interest were cloned. Whenever possible, we chose to validate genes that are the last in operons to avoid polar effects of CRISPRi. All spacers were cloned into the pWJ587 backbone with a chloramphenicol-resistant (cm^R^) marker.

Pairwise competition assay was performed to calculate the fitness of RN4220 carrying spacers targeting genes of interest (candidate), relative to RN4220 carrying no spacer (ie, pWJ700), which acted as the common competitor. First, candidate and the common competitor strains were streaked on TSA plates supplemented with 5 µg/mL chloramphenicol and incubated at 37°C for 14 – 20 hours. Equal amount of candidate the common competitor cells were mixed by picking and re-suspending single colonies in 1X PBS to an OD_600_ of 0.1 measured by plate reader. 2 uL of mixed bacteria were inoculated into 200 uL TSB containing 5 µg/mL chloramphenicol, IPTG, and testing antibiotics whenever necessary. Microplates were grown at 37 °C with shaking (548 cpm) for 20 – 24 h. After growth, 150 µL of bacteria in each condition were transferred to an 1.5 mL Eppendorf tube, pelleted and washed with 1X PBS. Importantly, as candidate genes were essential, it was necessary to choose IPTG concentrations that reduced the fitness of these genes by 50-90% relative to that of the common competitor under no antibiotic treatment.

To quantify the ratio of candidate and common competitor, pelleted cells were lysed, minipreped, and subjected to PCR. For each PCR, a 20 µL reaction mix was prepared by adding approximately 20 ng plasmid DNA as template, 0.2 uM forward primer (W1201), 0.2 uM reverse primers (L401) and One*Taq*® DNA polymerase (NEB). DNA bands were visualized on 2% agarose gel containing ethidium bromide. Intensity of DNA bands corresponding to spacers of interest (211 bp) and the common competitor (empty array, 145 bp) were quantified by ImageJ. An example of *ftsZ* is shown in **Fig. S1i**. At the beginning and end of pairwise competition assay, we also spotted bacteria on TSA plates containing 5 µg/mL chloramphenicol and incubated at 37°C for 14 – 20 hours. Combined with the ratios obtained from PCR, it allowed us to determine the initial and final population size of the candidate and the common competitor. Thus, fitness of candidate strain with spacer targeting gene *X* (W_X_) relative to that of the common competitor ^67,68^ can be calculated as:

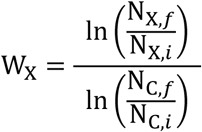

where N_X_ and N_C_ are the population sizes of gene *X* strain and the common competitor, and subscripts *f* and *i* indicate the final and initial time points, respectively.

We also performed a mock pairwise competition between RN4220 carrying pWJ587 (WT) and RN4220 carrying pWJ700 (the common competitor) under plain and various antibiotic conditions. Essentially, two WT strains were being competed. As expected, the relative fitness of WT cells under various antibiotic conditions were not significantly different from that in plain media (**Fig. S1j**).

### Checkerboard assays for measuring MIC

A single colony (10^6^-10^7^ CFUs) of *S. aureus* RN4220 cells carrying a chloramphenicol-resistant plasmid encoding IPTG-inducible dCas9 and a spacer targeting gene of interest were re-suspended in 100 µL 1X PBS. After serial dilution, approximately 50 – 200 CFUs were spotted on 5 x 4 TSA plates containing 5 µg/mL chloramphenicol (to keep the CRISPRi plasmid targeting genes of interest) and various concentrations of IPTG and testing antibiotics. The checkerboard format was necessary as a range of IPTG concentrations was needed to capture the optimal transcriptional repression levels for different essential genes. Typically, TSA was supplemented with following IPTG concentrations: 0.003, 0.01, 0.03, 0.1, and 0.3 mM.

To complement *ruvB*, RN4220 cells carrying dual plasmids pWJ658 (a chloramphenicol-resistant plasmid encoding IPTG-inducible dCas9 and a spacer targeting *ruvB*) and pWL15 (a tetracycline-resistant plasmid encoding IPTG-inducible *ruvB* containing synonymous mutations that negate CRISPRi targeting) were spotted on TSA plates containing 5 µg/mL chloramphenicol, 2.5 µg/mL tetracycline, appropriate amount of IPTG, and testing antibiotics. Similarly, *ligA* complementation was performed using RN4220 cells carrying plasmids pWJ653 and pWL25.

TSA plates were incubated at 37°C for 24 or 48 hours.

### Network analysis

To construct an essential gene similarity network in *S. aureus* RN4220 (**Figs. 5a** and **5b**), we performed our analysis on genes determined to be essential by either Santiago’s^23^ or Bae’s^25^ Tn-seq studies. Pairwise Pearson correlation coefficients (PCCs) between these essential genes were calculated based on their relative fitness in 14 antibiotic conditions (oxa 0.05, dapt 0.25, dapt 0.5, vanc 1, cipro 0.2, cipro 0.4, tmp 0.4, gent 0.5, gent 1, erm 0.08, lzd 0.2, lzd 0.25, tet 0.04, and mup 0.02), generating a 440 x 440 gene-gene correlation matrix (**Table S12**). This matrix was used to generate a gene similarity network, using the Kamada-Kawai algorithm. The threshold of PCC used was 0.65. To gain process-level insights, the mean of pairwise PCCs of essential genes from any two biological processes were calculated and subjected to hierarchical clustering (**Fig. 5c**).

### Gene-gene interaction measurement

To measure genetic interactions, *S. aureus* RN4220 carrying dual-plasmid systems were cloned. A chloramphenicol-resistant plasmid driving a xylose-inducible antisense RNA fragment was used to repress CC genes. Another tetracycline-resistant plasmid driving an IPTG-inducible dCas9 and crRNA was used to repress DD genes, *coaA*, and *ribF*. For each dual-gene combination, 5 – 10 single colonies of *S. aureus* RN4220 carrying dual-plasmid systems were re-suspended in 1X PBS. Serial dilutions were prepared and appropriate numbers of CFUs were plated on TSA containing 5 µg/mL chloramphenicol, 2.5 µg/mL tetracycline, and various combinations of IPTG and xylose. TSA plates were incubated for 22 hours at 37 °C, after which colony areas were quantified using ImageJ as a proxy for bacterial fitness. For each gene, we selected inducer concentrations that reduced fitness to 20%-90% of unperturbed cells for subsequent quantification. For each gene pair (gene A and gene B), the fitness of individual gene perturbations and dual gene perturbation were normalized to that of cells with no perturbation, generating W_A_, W_B_, and W_A, B_, respectively. Genetic interaction, or epistasis, was calculated as ε_A,B_ = W_A, B_ - W_A_ • W_B_. As controls, the fitness of bacteria carrying a non-targeting two-plasmid system (ie, parental plasmids pC194 and pWL17) in the presence of various combinations of IPTG and xylose were also measured, allowing for the calculation of baseline epistasis. To determine the statistical significance of genetic interactions, a two-sample t-test between triplicates of test gene-gene pairs (ε_A,B_) and triplicates of non-targeting two-plasmid system induced in the same IPTG and xylose concentrations (ε_pC194,pWL17_) were performed.

### RNA extraction, reverse transcription, and quantitative PCR (qPCR)

*S. aureus* RN4220 strains harboring IPTG-inducible CRISPRi system targeting DD genes or null control were grown overnight on TSA plates containing 5 µg/mL tetracycline at 37 °C. The next day, approximately 100 colonies were pooled and inoculated into fresh TSB supplemented with 1 mM IPTG to induce CRISPRi repression of target gene, followed by incubation with shaking (220 rpm, 37 °C) for 1 h. Approximately 10 units (1 unit = 1 mL bacterial culture at OD_600_ of 1) of cells were harvested by centrifugation at 5,000 × g for 5 min at room temperature, resuspended in RNAlater™ Stabilization Solution (Invitrogen), incubated at room temperature for 30 min, and stored at 4 °C until RNA extraction.

RNAlater-treated cell pellets were transferred into a 2-mL bead-beating tube preloaded with 0.1-mm glass beads. Cells were resuspended in 500 µL Buffer RLT (RNeasy Mini Kit, Qiagen) freshly supplemented with 1% β-mercaptoethanol (10 µL per mL buffer) and vortexed briefly (5–10 s). Tubes were processed in a bead beater at 30.0 m/s for 30 s, four times, with samples kept on ice for 30 s between cycles to prevent overheating. Lysates were clarified by centrifugation at 13,000 × g for 1 min at 4 °C, and the supernatants were carefully transferred to fresh RNase-free tubes without disturbing beads or debris. The clarified lysates were then subjected to RNA purification using the RNeasy Mini Kit (Qiagen), followed by in-solution DNase I treatment to remove genomic DNA. RNA integrity was assessed using a TapeStation system (Agilent), and only samples with RIN > 8 were used for downstream analyses.

cDNA was synthesized from 500 ng – 1 µg total RNA using Maxima H Minus Reverse Transcriptase (Thermo Scientific) with random hexamer primers, according to the manufacturer’s protocol. To quantify the expression of *sosA* (SAOUHSC_01334), qPCR was performed using primers WL83 and WL84 and Power SYBR™ Green PCR Master Mix (Applied Biosystems) on a QuantStudio 6 Real-Time PCR System (Applied Biosystems). Primers W1643 and W1644 were used to amplify the housekeeping *gmk* (SAOUHSC_01176). For each sample, relative expression of *sosA* was calculated using ΔCt=Ct_sosA_−Ct_gmk_. The fold change of *sosA* expression in experimental strain compared to control unperturbed strain was calculated as 2^−ΔΔCt^, where ΔΔCt=ΔCt_experimental_−ΔCt_control_. All primers were validated for efficiency and linear range of amplification using standard qPCR approaches. Specificity was confirmed through melting curve analysis.

### Checkerboard assays for measuring antibiotic-SMI interactions

Synergy H1 microplate reader (Biotek) was used to perform checkerboard assays. Typically, Corning™ 3370 Clear Polystyrene 96-Well Microplates were used to harbor a 7 x 7 checkerboard containing 200 µL TSB with varying concentrations of antibiotics and SMIs. To demonstrate the potentiation of antibiotics by SMIs, it was necessary to use the concentrations of antibiotics, but not those of the SMIs, in 2-fold serial dilutions. 5 – 10 single colonies of *S. aureus* RN4220 or USA300 cells were re-suspended in 200 µL 1X PBS. 2 µL of re-suspended cells were inoculated into 200 µL media in each well of the checkerboard. Microplates were grown at 37°C with shaking (548 cpm) for 22 – 24 h. The absorbance at 600nm (OD_600_) was measured every 10 minutes.

We use **Fig. S10a** as an example to illustrate how antibiotic-SMI interactions are quantified. First, for each time point, bacterial growth in various concentrations of antibiotic and SMI was normalized to bacterial growth in plain media (ie, 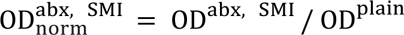). Panel (i) shows normalized bacterial growth at the terminal time point (22-24 hours post-inoculation). An asterisk “*” labels the well where the change in MIC (ΔMIC) was determined. However, it is well known that terminal-point MIC determination can sometimes overestimate drug-drug interactions. To be more stringent, we adopted an additional epistasis metric from a recent large-scale drug-drug interaction study in bacteria^50^, with modifications. Panel (ii) shows the four growth curves related to the well labeled with asterisk: plain media (gray), antibiotic alone (black), SMI alone (blue), and both antibiotic and SMI combined (red). For each time point along these growth curves, epistasis between antibiotic and SMI (ε^abx, SMI^) was calculated using a Bliss interaction model: 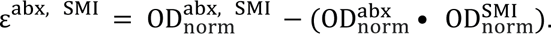 For the well where ΔMIC was determined (ie, “*”), we calculated ε^abx, SMI^ at 1-hour intervals, from 6 hours (when bacteria in plain media entered stationary phase) to 24 hours, the terminal time point. These values were averaged to yield the final ε, which quantifies the level of antibiotic-SMI interaction.

By quantifying ε^abx, SMI^ over an extended period of bacterial growth, the epistasis metric effectively compliments the biased MIC determination based on a single time point. For example, D-pantothenol and cipro seemingly synergized as D-pantothenol reduced the MIC of cipro by 4-fold in the 24-hour checkerboard result (**Fig. S10l**). However, this proved to be an overestimation, as growth curves clearly revealed that bacteria only began growing in D-pantothenol (blue curve) at a late time point. Epistasis measurements showed that the average ε^abx, SMI^ from 6 to 24 hours was -0.06, indicating that the synergy was rather weak. We only considered a simultaneous reduction in MIC and a strong negative epistasis (ΔMIC ≤ ½ and ε ≤ -0.4) to be a synergistic antibiotic-SMI interaction.

## FIGURES AND LEGENDS

**Fig. S1.**
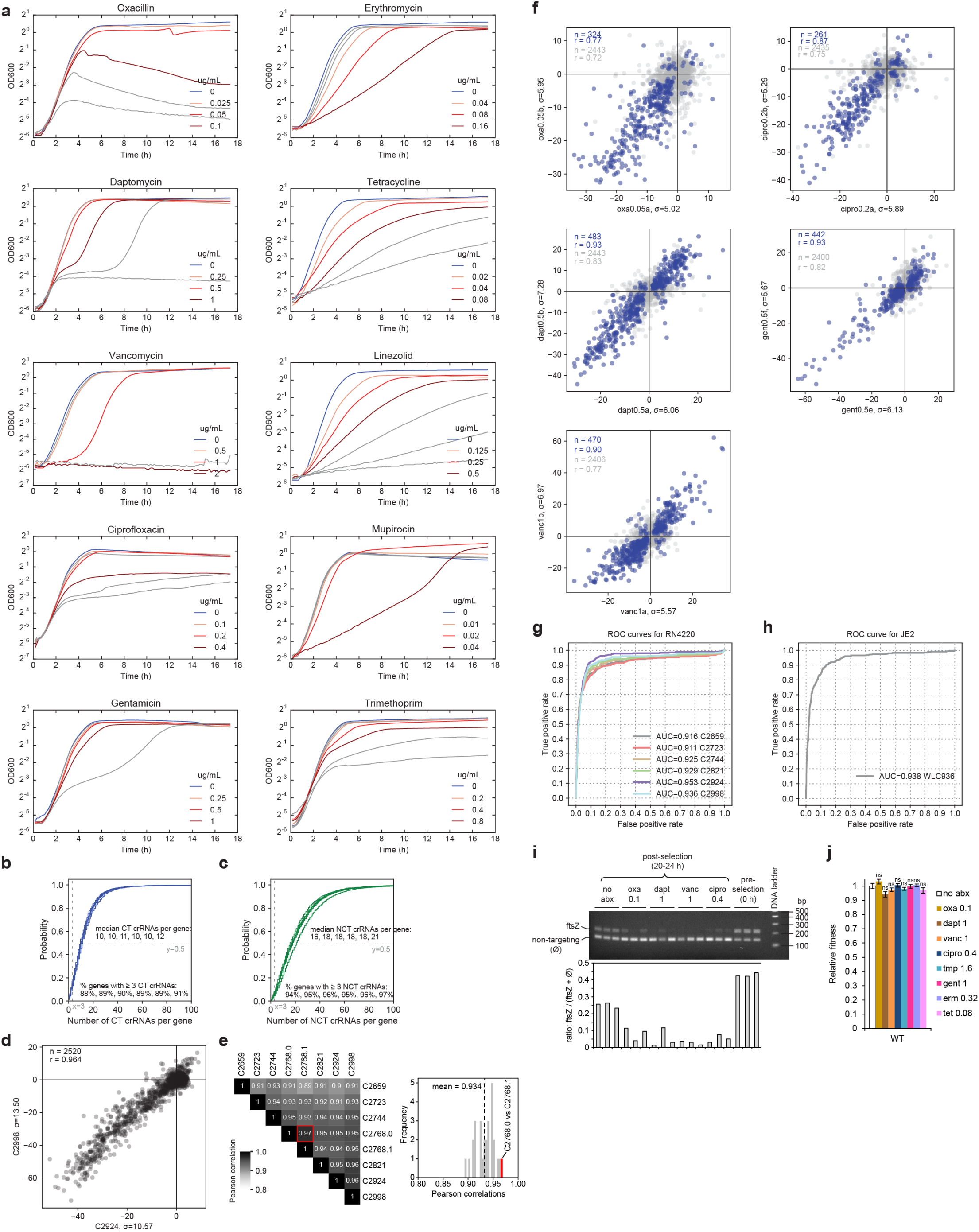
Antibiotic conditions and quality controls. (**a**) Titration growth curves of *S. aureus* RN4220 grown in ten antibiotics used in this study. Concentrations most relevant to the CRISPRi screens are shown in warm colors. (**b**) Cumulative distribution function plot of the number of coding-strand-targeting (CT) crRNAs per gene from six independent CALM-CRISPRi libraries made in RN4220. Libraries created by CALM-CRISPRi are highly comprehensive, with 88-91% of genes covered by 3 or more CT crRNAs, and a median CT crRNAs per gene being 10-12 from these replicates. (**c**) Same as (b) except plot is showing the noncoding-strand-targeting (NCT) crRNAs. (**d**) Pearson correlation between the fitness scores of RN4220 genes (Z_Ø_) from two independent CALM-CRISPRi libraries grown in TSB for 9 hours. Only genes targeted by at least 2 CT crRNAs were included in analysis. (**e**) Left: Pairwise Pearson correlations of the fitness scores of RN4220 genes (Z_Ø_) from all seven independent CALM-CRISPRi libraries grown in TSB for 9 hours. Libraries C2768.0 and C2768.1 were made in the same experiment, serving as a batch control (red box) for other libraries generated in separate experiments. Right: Distribution of all Pairwise Pearson correlations on the left panel. Red bar indicates correlation between C2768.0 and C2768.1. (**f**) Pairwise Pearson correlations between the relative fitness scores of RN4220 genes in antibiotics (Z_abx_) from two independent CALM-CRISPRi libraries. The numbers of genes passing quality control and their correlations are shown in gray in upper left corners. The numbers of genes with significantly altered fitness in antibiotics relative to TSB (|Z_abx_| ≥ 9, P_adj_ < 0.05) and their correlations are shown in blue. (**g**) ROC curves of six independent CALM-CRISPRi libraries generated in *S. aureus* RN4220 and grown in TSB for 9 hours. Gene essentiality determined by Santiago’s Tn-seq study^23^ was used as a reference. (**h**) ROC curve of CALM-CRISPRi library generated in *S. aureus* JE2 and grown in TSB for 9 hours. Gene essentiality determined by Coe’s Tn-seq study^4^ was used as a reference. (**i**) An example of the quantification of pairwise competition assays in triplicates. Equal amount of RN4220 with a spacer targeting *ftsZ*, and RN4220 with a non-targeting spacer (ie, the common competitor, Ø) were mixed, representing the pre-selected samples at 0 h. Mixed bacteria were inoculated into TSB and TSB containing various antibiotics (concentration shown as µg/mL) and grown for 20 - 24 hours at 37 °C. DNA from pre- and post-selected samples were used as templates for PCR amplifying the CRISPR spacer region and visualized on agarose gel (upper panel). The ratios of normalized intensity of *ftsZ* to total (*ftsZ* + Ø) were calculated and shown as bar plots (lower panel). (**j**) Pairwise competition assays (Methods) measuring the relative fitness of *S. aureus* RN4220 carrying an IPTG-inducible CRISPRi system that does not have a genomic target (ie, WT) in indicated antibiotic conditions. Essentially, two WT strains were being competed. Concentrations of antibiotics are shown in µg/mL. Error bars indicate the standard deviation from three biological replicates. For each gene knockdown, paired t-tests were performed between the antibiotic condition and the no-antibiotic condition.

**Fig. S2.**
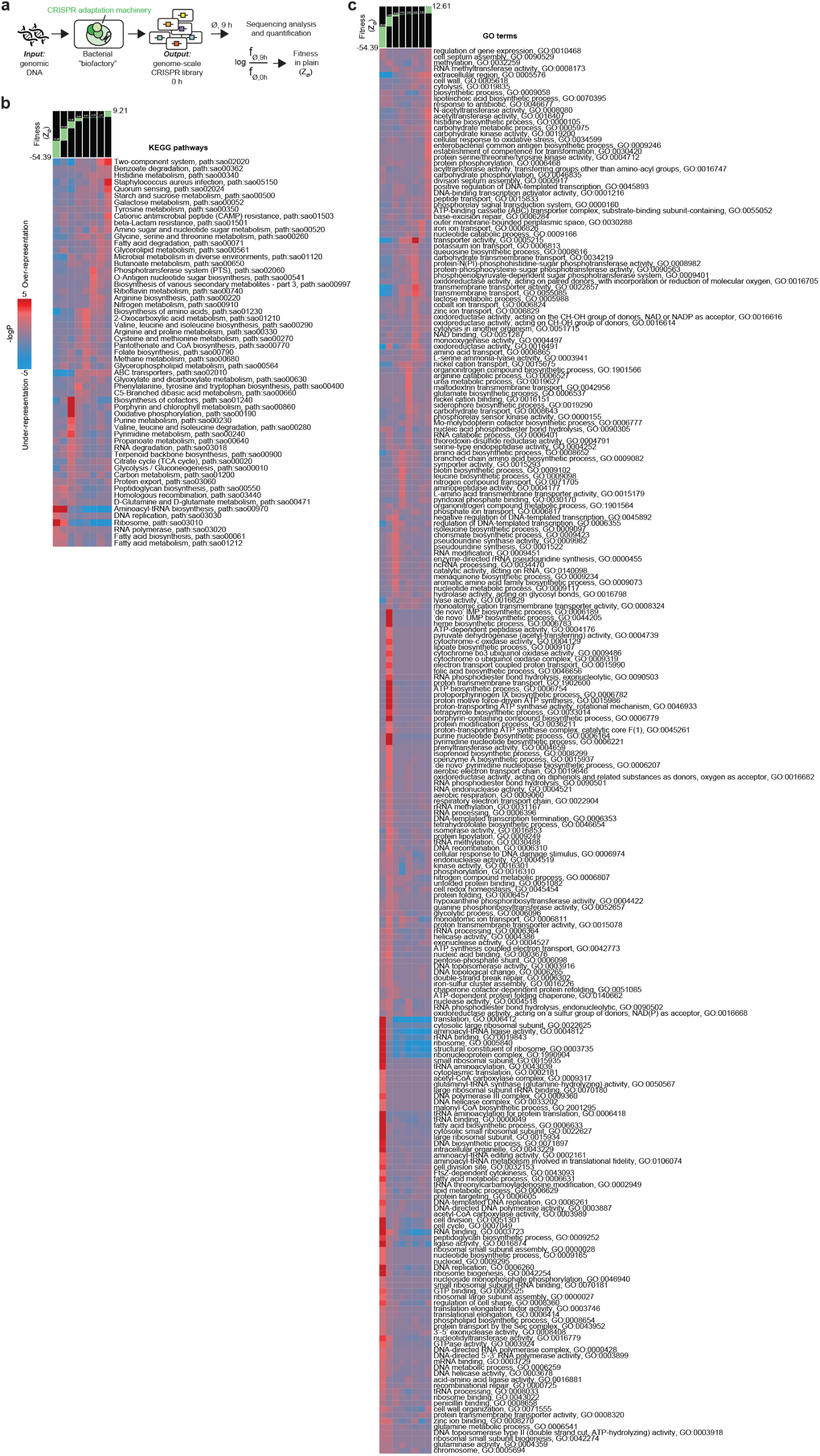
Functional enrichment analysis of the gene essentiality landscape in *S. aureus* RN4220. (**a**) CALM-CRISPRi libraries of *S. aureus* RN4220 were grown in plain media, TSB, for 9 hours. Gene fitness in TSB (Z_Ø_) was quantified by sequencing analysis (Methods). For each gene, the mean Z_Ø_ was calculated from three biological replicates. (**b**) KEGG pathway enrichment analysis was performed using the mean Z_Ø_ values of genes and the iPAGE tool^60^. Z_Ø_ of genes was ranked and binned. For each bin, the top black-green panel shows the lower and upper bounds of Z_Ø_, and the bottom red-blue panel shows the degree of overrepresentation (red) and underrepresentation (blue) of genes belonging to the KEGG pathway. (**c**) Same as (b) except iPAGE enrichment analysis was done on GO terms.

**Fig. S3.**
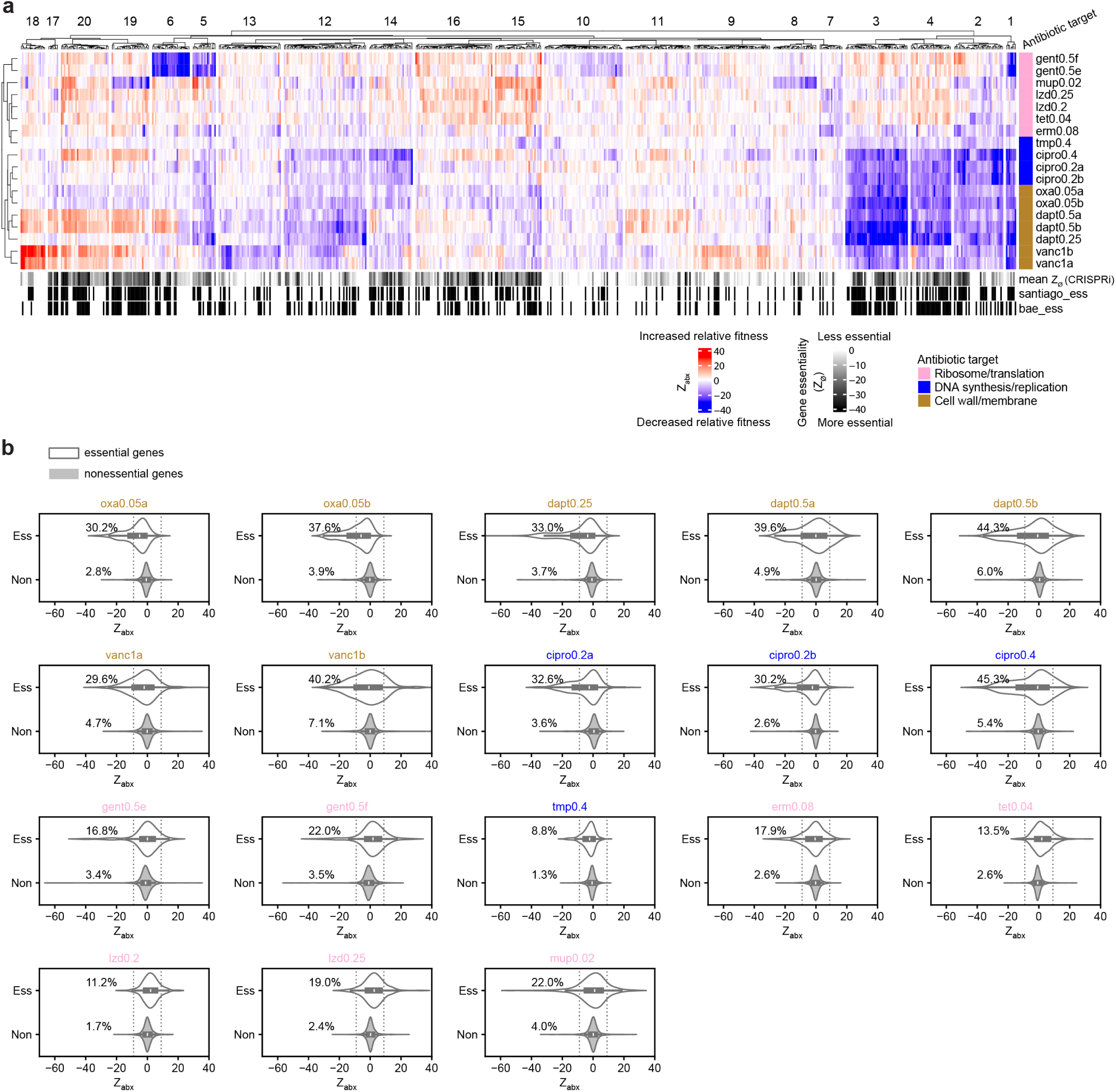
Antibiotic-gene interactions in *S. aureus* RN4220. (**a**) Hierarchical clustering of 650 genes whose repression by CALM-CRISPRi significantly increased (red) or decreased (blue) relative fitness in at least one antibiotic condition (|Z_abx_| ≥ 9 and P_adj_ < 0.05). A total of 18 antibiotic conditions (including biological replicates) is shown. For each gene, its essentiality in plain TSB (black and white) was quantified as Z_Ø_ by CRISPRi (mean of triplicates) in this study, and qualified (binary) by Bae’s^25^ and Santiago’s^23^ Tn-seq studies. Antibiotics are color coded by their targets. Z_Ø_, Z_abx_, and clustering are shown in Tables S2, S3, and S5, respectively. (**b**) Distributions of Z_abx_ of essential and nonessential genes in 18 antibiotic conditions. In each panel, dotted lines indicate |Z_abx_| = 9, and percentage of genes with |Z_abx_| ≥ 9 within essential (ess) and nonessential (non) genes are shown.

**Fig. S4.**
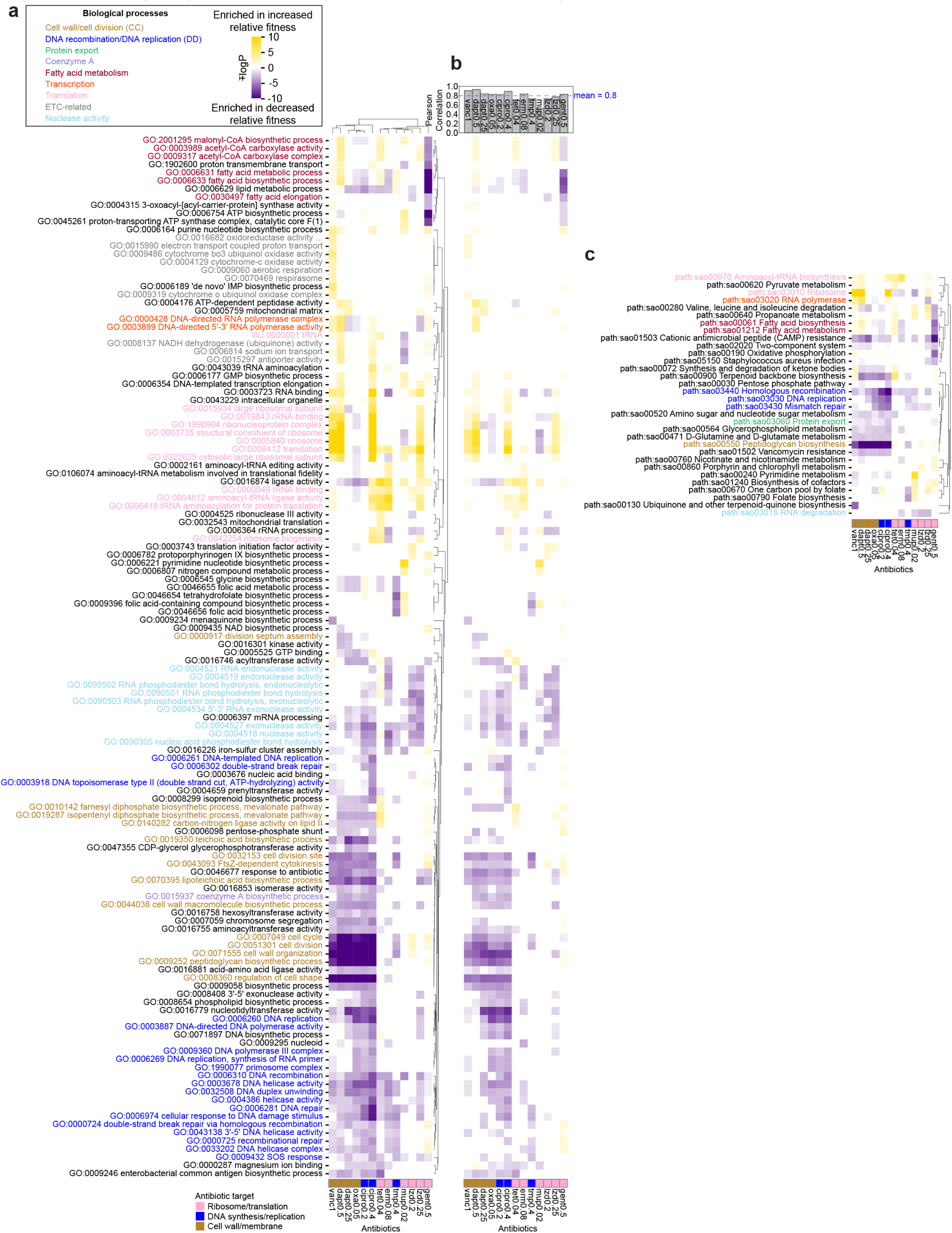
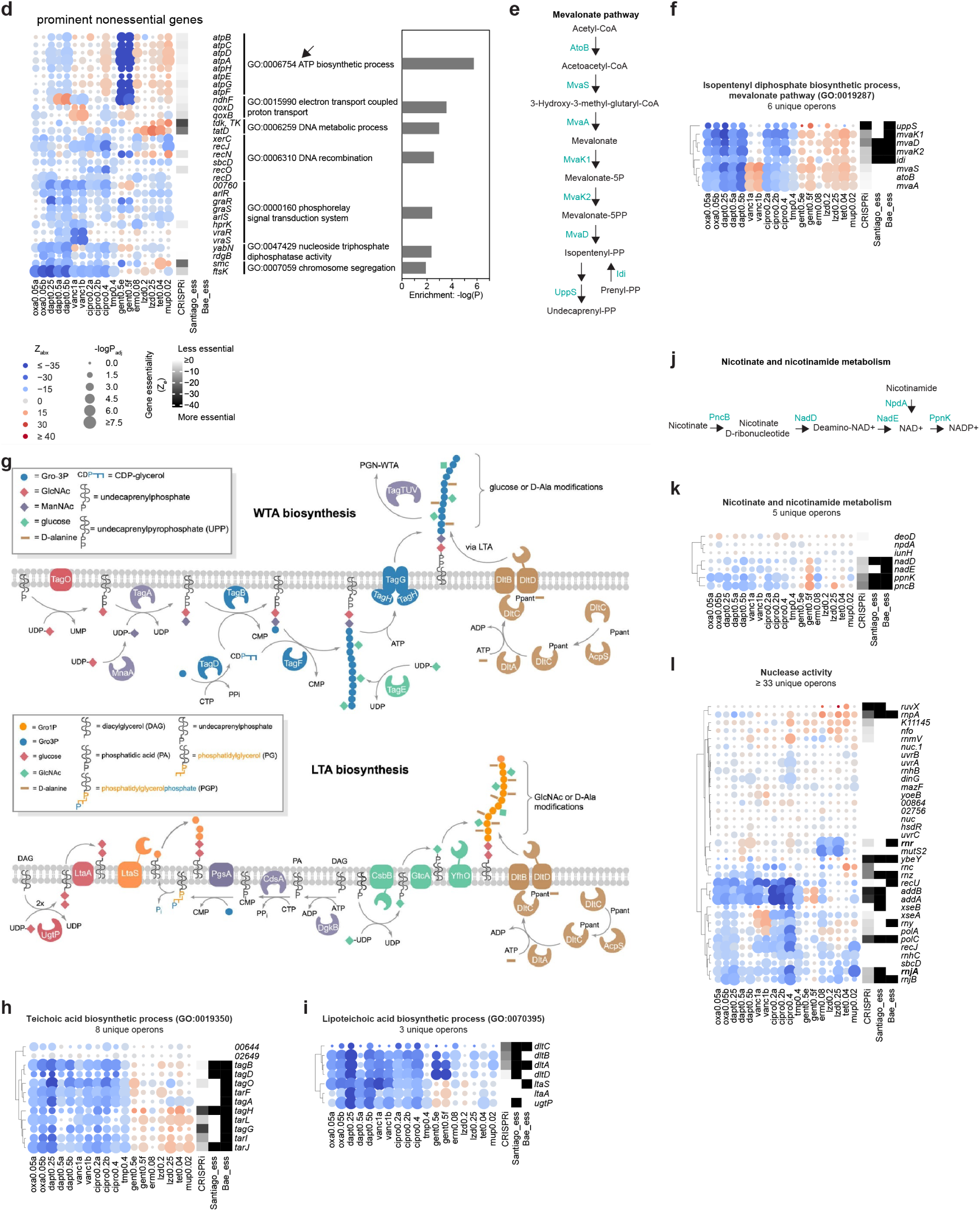
Biological processes that modulate antibiotic sensitivity in *S. aureus* RN4220. (**a**) Functional enrichment analysis by iPAGE^60^ followed by hierarchical clustering of all significantly enriched GO terms. The ∓log(P-value) of GO terms significantly enriched (P < 0.001) for genes whose repression increased and decreased relative fitness (|Z_abx_| ≥ 9 and P_adj_ < 0.05) are shown in yellow and purple, respectively. Color code for biological processes is the same as Fig. 1g. We added a new term called “WJ:0000001 tRNA” as tRNA genes are annotated in neither GO terms nor KEGG pathways. (**b**) Bottom: Same as (a) except that enrichment analysis was performed using only the last gene in each of the 1691 operons annotated by BioCyc (http://biocyc.org/). Rows and columns are in the same order as those in (a). Clustering of all GO terms is shown in Table S7. Top: Pearson correlation coefficient for each antibiotic condition between enrichment analyses performed using full gene set and the last gene in each operon. (**c**) Same as (a) except showing clustering of all significantly enriched KEGG pathways. Antibiotic columns are in the same order as those in (a). (**d**) Biological processes that were enriched for nonessential genes whose repression significantly altered relative fitness in antibiotics. Left: heatmap showing the relative fitness of genes in ten antibiotics as quantified by CALM-CRISPRi screens (Z_abx_). Gene names are annotated by KEGG orthology (ko). Gene knockdowns that decreased and increased relative fitness in antibiotics are shown in blue and red circles, respectively. Size of circle indicates the negative of log-transformed adjusted P-value, -logP_adj_. Gene essentiality quantified by CRISPRi (mean of Z_Ø_ from triplicates) and qualified (binary) by Santiago’s^23^ and Bae’s^25^ Tn-seq studies are also shown. Right: name of biological processes and their level of enrichment shown as -log(P). (**e**) Schematic of the mevalonate pathway. (**f**) Same as the left panel of (d) except heatmap is showing genes in the isopentenyl diphosphate biosynthetic process, mevalonate pathway (GO:0019287). *atoB*, *mvaS*, *mvaA*, *idi* and *uppS* were also added to the heatmap due to overannotation by GO terms. (**g**) Wall teichoic acid (WTA) biosynthesis and lipoteichoic acid (LTA) biosynthesis pathways in *B. subtilis*. Schematic is adapted from Walter et al^61^ with minor modifications. (**h**) Same as the left panel of (d) except heatmap is showing genes in teichoic acid biosynthetic process (GO:0019350). *tagGHO* were also added to the heatmap due to overannotation by GO terms. (**i**) Same as the left panel of (d) except heatmap is showing genes in lipoteichoic acid biosynthetic process (GO:0070395). (**j**) Schematic of nicotinate and nicotinamide metabolism. (**k**) Same as the left panel of (d) except heatmap is showing genes in nicotinate and nicotinamide metabolism (KEGG path:sao00760). (**l**) Same as the left panel of (d) except heatmap is showing genes in the nuclease activity (GO:0004518).

**Fig. S5.**
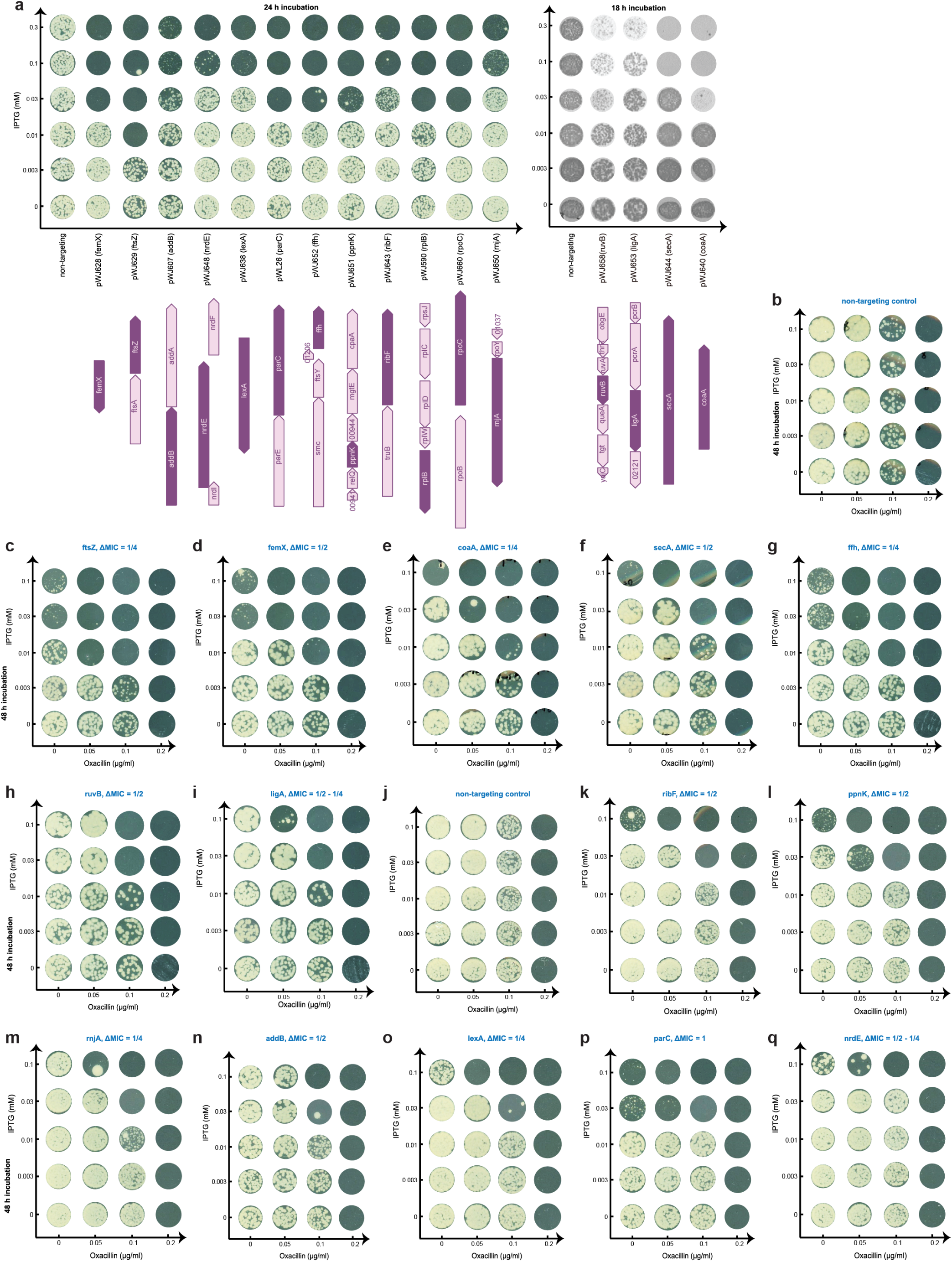
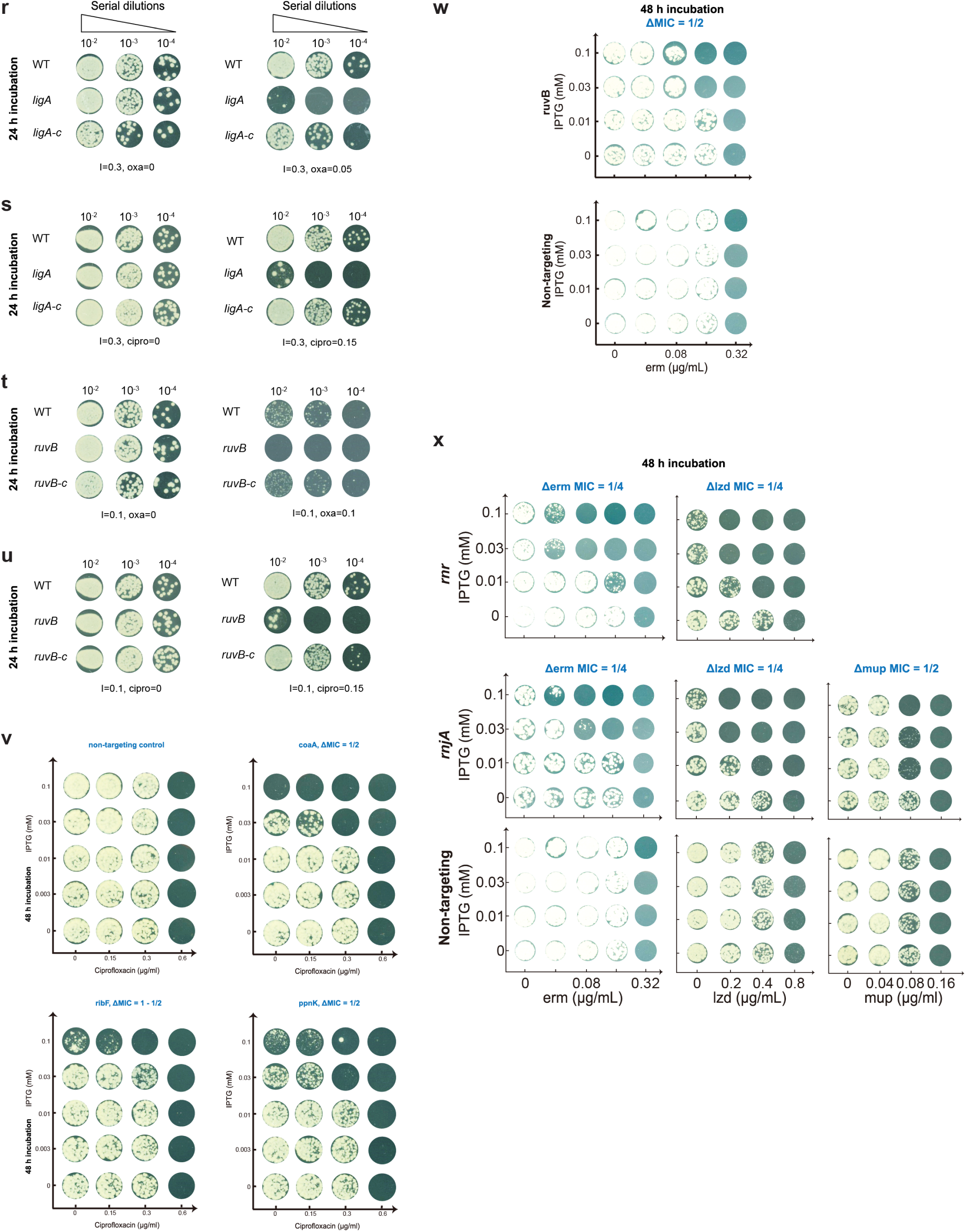
Gene essentiality and checkerboard MIC assays on solid agar. (**a**) *S. aureus* RN4220 carrying an IPTG-inducible CRISPRi system targeting various essential genes or a non-targeting CRISPRi system grown on TSA supplemented with varying concentrations of IPTG. Approximately 50 – 200 CFUs were spotted on these plates and grown for 24 or 18 hours at 37 °C. Genes of interest and other genes in their operon, if present, are shown in plum and pink, respectively. (**b-i**) Checkerboard MIC assays of oxacillin for *S. aureus* RN4220 carrying an IPTG-inducible CRISPRi system targeting seven essential genes and a non-targeting control. For each essential gene, it was necessary to perform a 5 x 4 checkerboard assay with varying IPTG concentrations on one axis in order to capture the optimal transcriptional repression level. Approximately 50 – 200 CFUs were spotted on TSA plates and grown for 48 hours at 37 °C. Change in MIC (ΔMIC) is shown in blue. (**j-q**) Same as (b-i) except checkerboard MIC assays with CRISPRi targeting another seven essential genes were done on another batch. (**r**) *ligA* complementation. “*ligA*” represents *S. aureus* RN4220 carrying a plasmid encoding an IPTG-inducible CRISPRi system targeting *ligA*. “*ligA-c*” represents *S. aureus* RN4220 carrying a plasmid encoding an IPTG-inducible CRISPRi system targeting *ligA*, complemented with another plasmid encoding an IPTG-inducible *ligA* containing synonymous mutations that negate crRNA-target base-pairing. Left: cells grown on 0.3 mM IPTG and 0 μg/mL oxacillin. Right: cells grown on 0.3 mM IPTG and 0.05 μg/mL oxacillin. Cells were grown on TSA plates for 24 hours at 37 °C and three dilutions were shown. (**s**) Same as (r) except antibiotic is ciprofloxacin. (**t**) Same as (r) except complementation was performed on *ruvB*. (**u**) Same as (t) except antibiotic is ciprofloxacin. (**v**) Same as (b-i) except that checkerboard assays measured MICs of ciprofloxacin for *S. aureus* RN4220 carrying an IPTG-inducible CRISPRi system targeting *coaA*, *ribF*, *ppnK* and a non-targeting control. (**w**) Same as (b) except that checkerboard assays measured MICs of erythromycin for *S. aureus* RN4220 carrying an IPTG-inducible CRISPRi system targeting *ruvB* and a non-targeting control. (**x**) Checkerboard assays measuring the MICs of erythromycin, linezolid, and mupirocin for *S. aureus* RN4220 carrying CRISPRi targeting *rnr* and *rnjA*. Approximately 50 – 200 CFUs were spotted on TSA plates and grown for 48 hours at 37 °C.

**Fig. S6.**
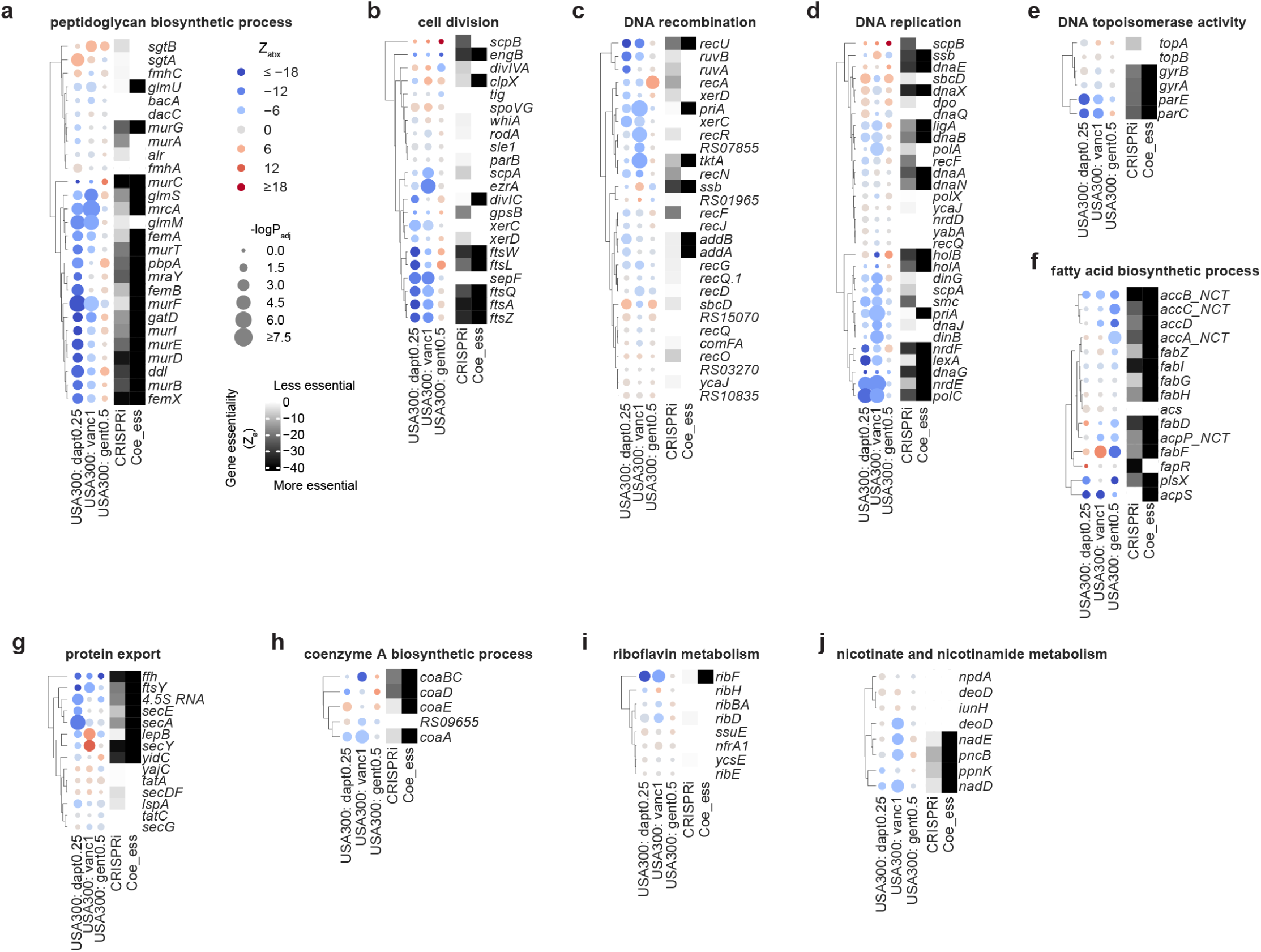
Select fitness profiles of methicillin-resistant *S. aureus* JE2 in antibiotics. (**a**) Heatmap showing the relative fitness of genes in the peptidoglycan biosynthetic process (GO:0009252) in three antibiotics as quantified by CALM-CRISPRi screens (Z_abx_). Gene names are annotated by KEGG orthology (ko). Gene knockdowns that decreased and increased relative fitness in antibiotics are shown in blue and red circles, respectively. Size of circle indicates the negative of log-transformed adjusted P-value, -logP_adj_. Gene essentiality quantified by CRISPRi (as Z_Ø_ in this study) and qualified (binary) by Coe’s Tn-seq study^4^ are also shown. (**b-j**) Same as (a) except heatmaps are showing genes in other biological processes. Z_Ø_ and Z_abx_ of all JE2 genes are shown in Tables S9 and S10, respectively.

**Fig. S7.**
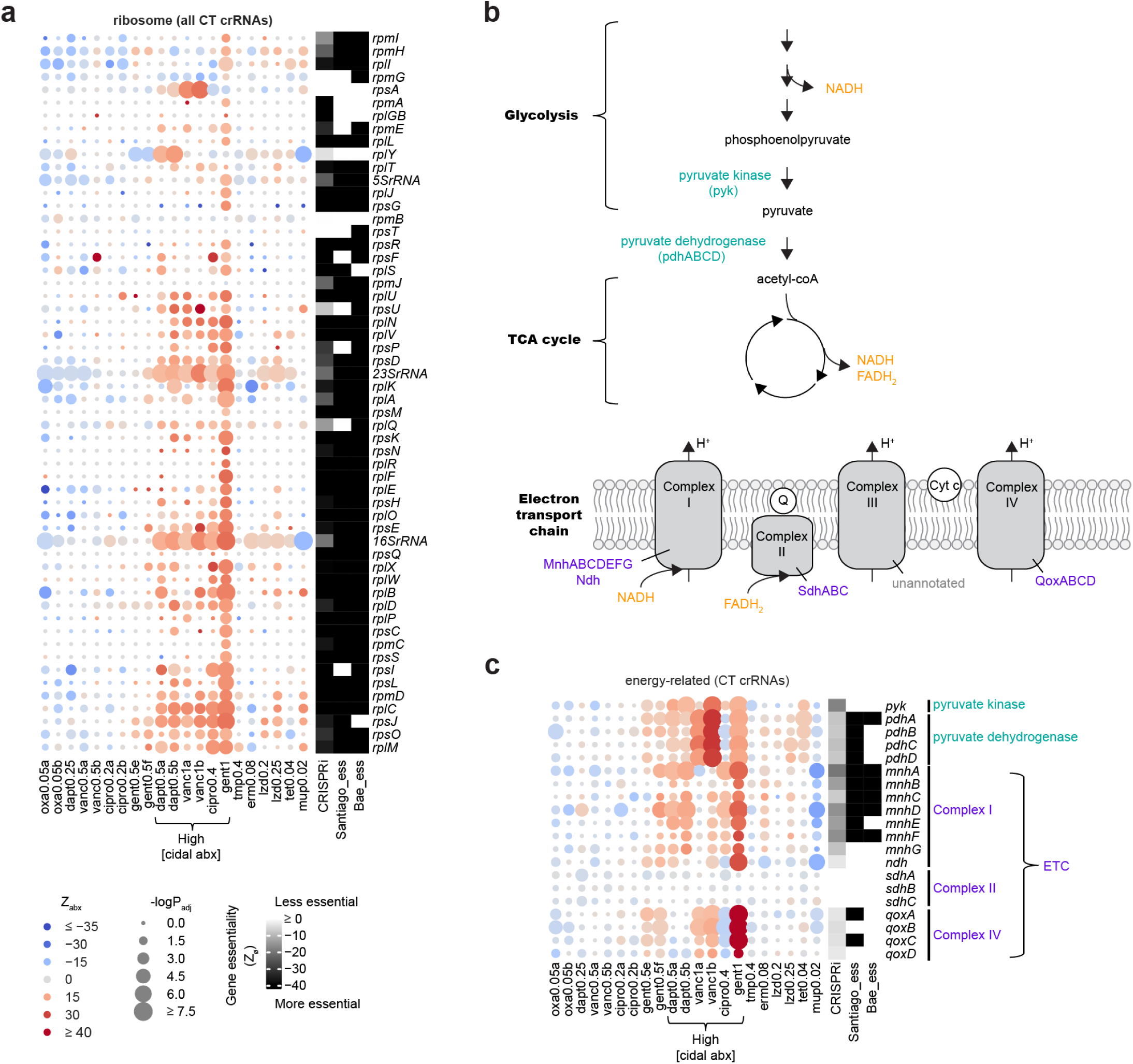
Other desensitizing antibiotic-gene interactions. (**a**) Heatmap showing the relative fitness of genes in ribosome (GO:0005840) in ten antibiotics as quantified by all CT crRNAs in CALM-CRISPRi screens (Z_abx_) in *S. aureus* RN4220. This quantification is incomplete (see “Caveat” in Methods). Gene names are annotated by KEGG orthology (ko). Gene knockdowns that decreased and increased relative fitness in antibiotics are shown in blue and red circles, respectively. Size of circle indicates the negative of log-transformed adjusted P-value, -logP_adj_. Gene essentiality quantified by CRISPRi (mean of Z_Ø_ from triplicates) and qualified (binary) by Santiago’s^23^ and Bae’s^25^ Tn-seq studies are also shown. (**b**) Simplified schematic of the glycolysis, TCA cycle and the electron transport chain. (**c**) Same as (a) except heatmap is showing the relative fitness of select genes in energy-related pathways shown in (b).

**Fig. S8.**
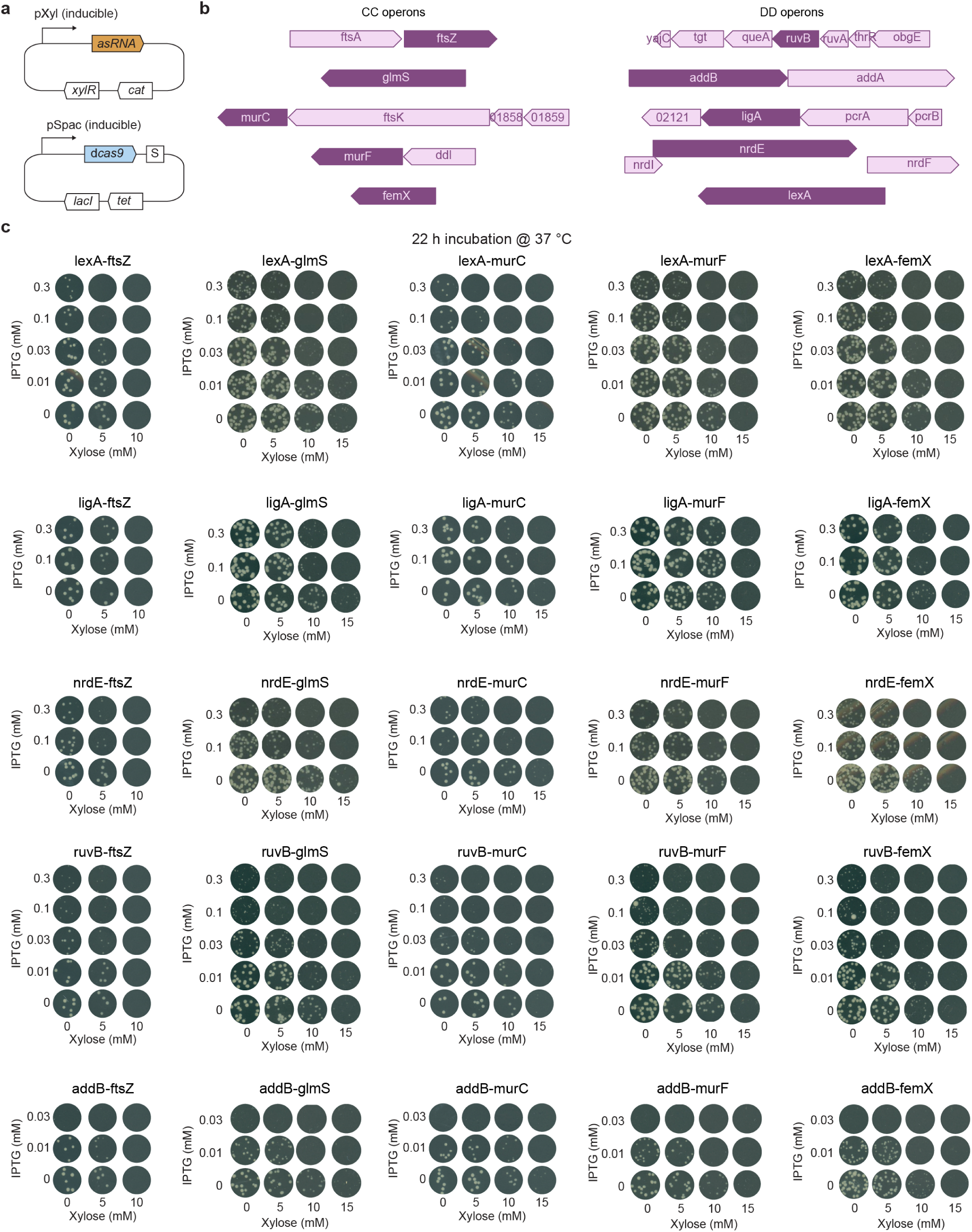
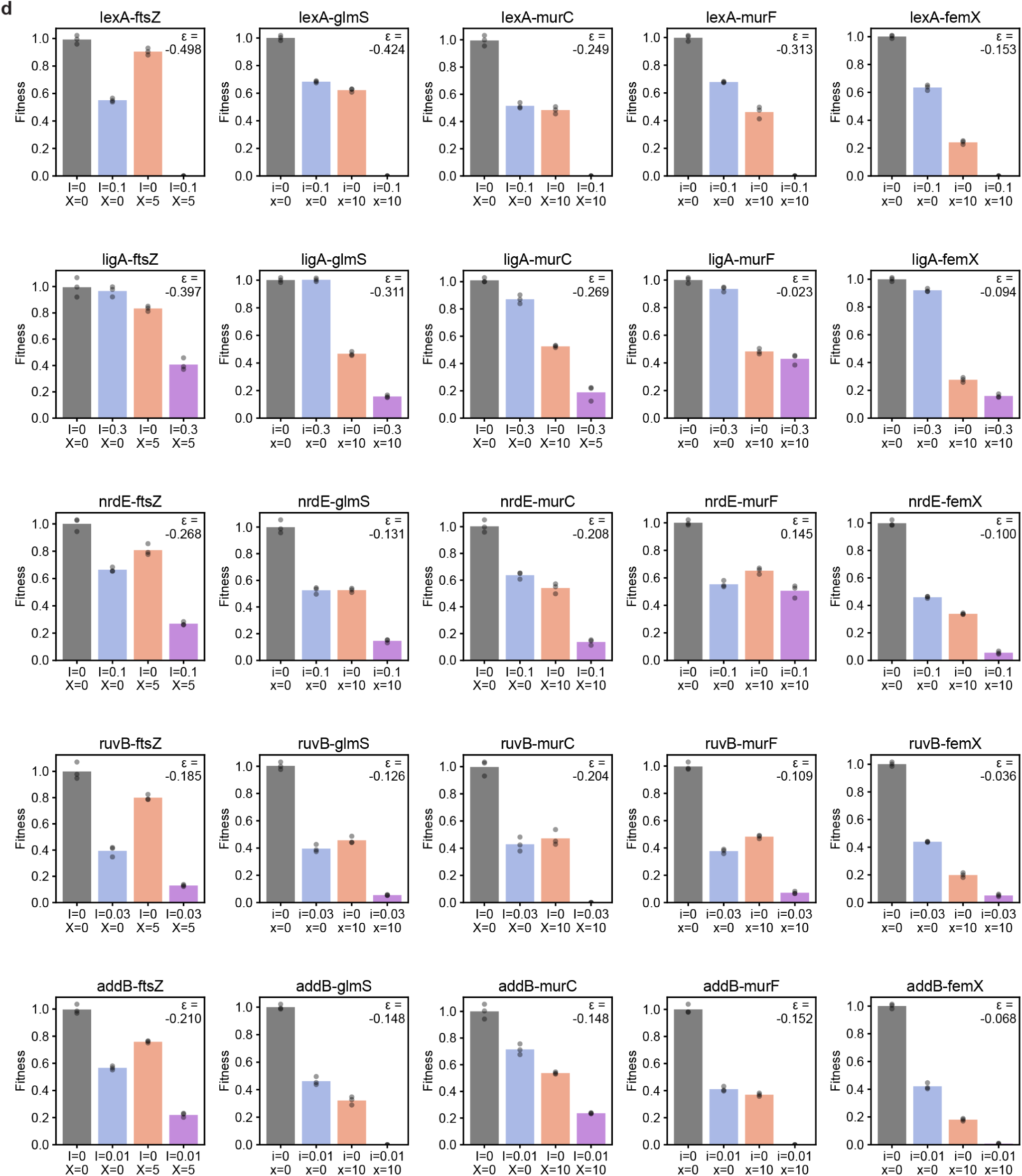
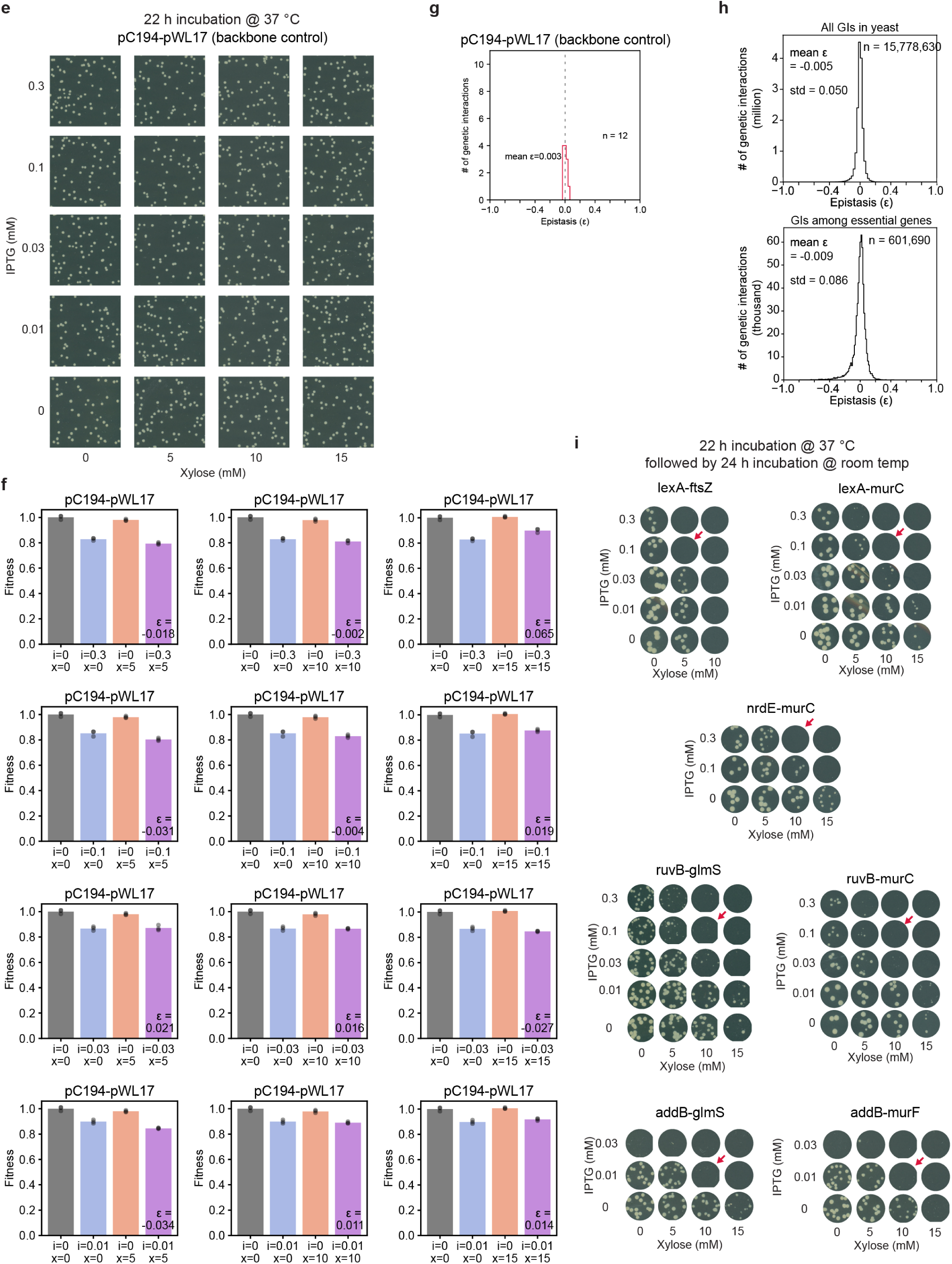
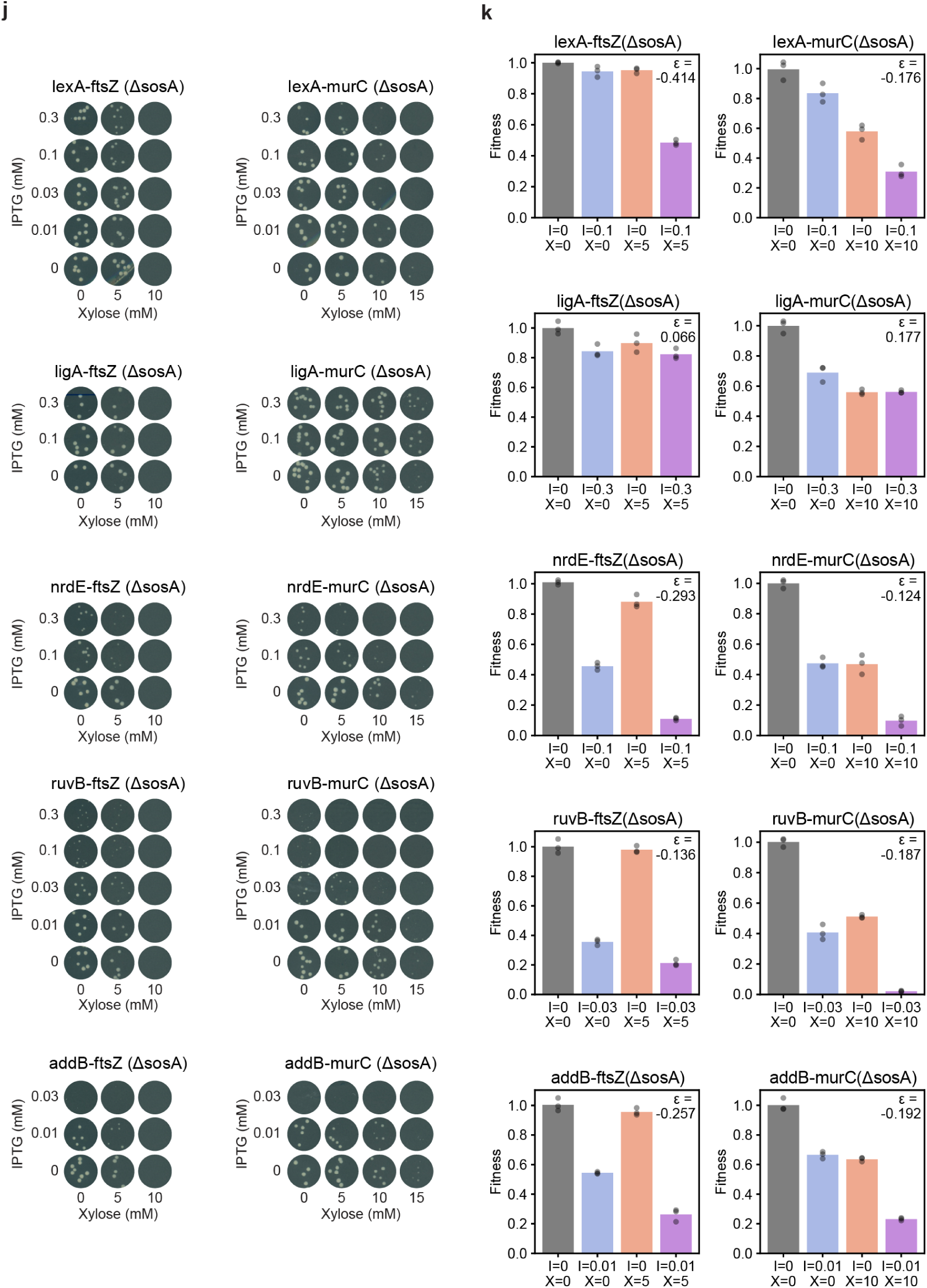
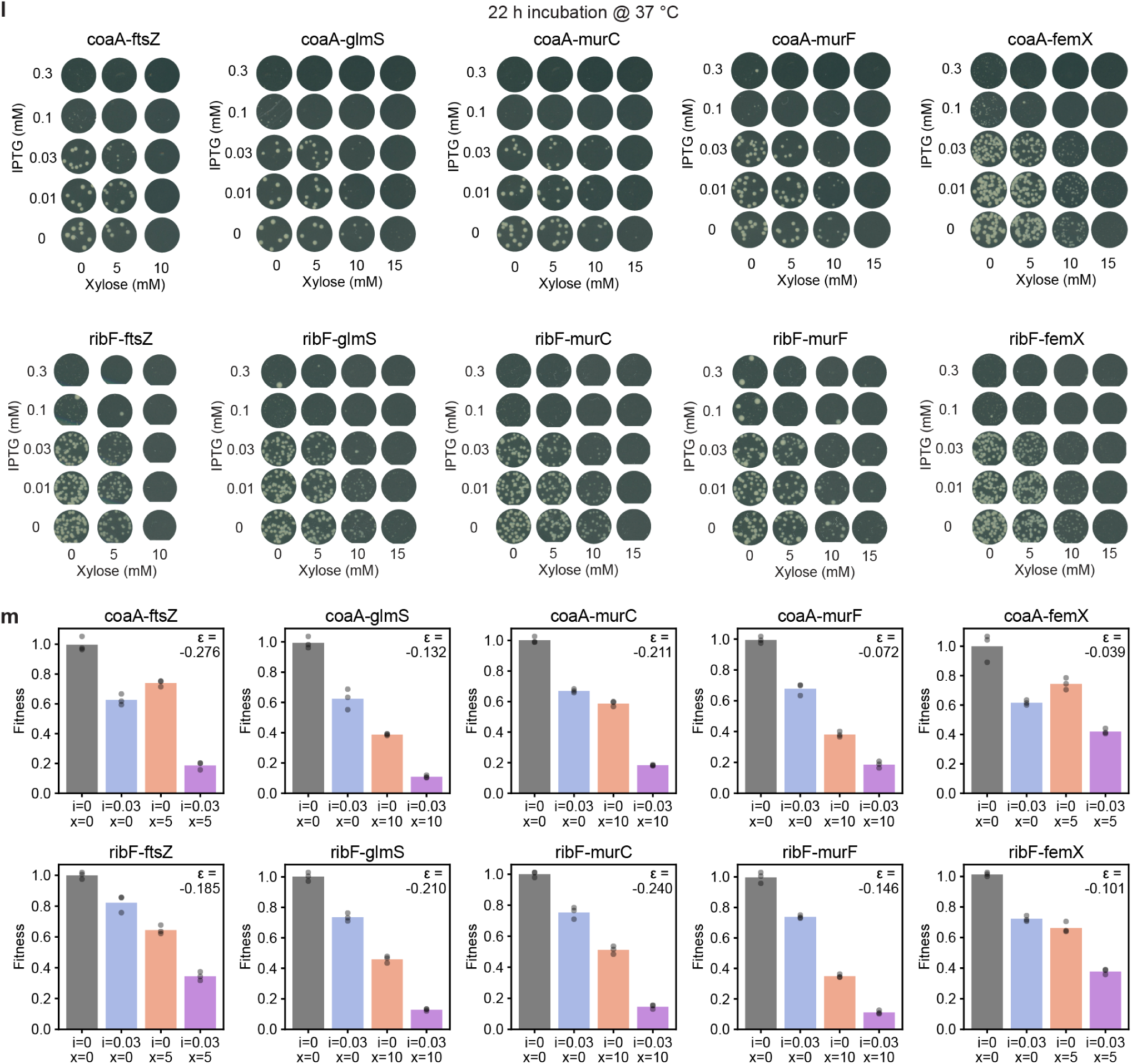
Quantification of genetic interactions. (**a**) The two-plasmid system used to quantify gene-gene interactions. pSpac and pXyl are IPTG- and xylose-inducible promoters driving the expression of dCas9/spacer (denoted as “S”) and antisense RNA (asRNA), respectively. Spacer is the DNA precursor to crRNA. (**b**) Operons of five CC and five DD genes. Targeted genes and other genes in their operons (if present) are shown in plum and pink, respectively. (**c**) Checkerboard assays to quantify genetic interactions between five CC genes and five DD genes. In each panel, RN4220 cells carrying IPTG-inducible dCas9 targeting a DD gene and xylose-inducible asRNA targeting a CC gene were plated on TSA plates containing various combinations of IPTG and xylose and grown for 22 hours at 37 °C. The order of genetic pairs is the same as those shown in Fig. 6a. (**d**) Quantification of fitness and epistasis for checkerboard assays in (c). Fitness was quantified by selecting appropriate inducer concentrations and measuring colony sizes (Methods). Epistasis between gene A and gene B (ε_A,B_) was calculated as ε_A,B_ = W_A, B_ - W_A_• W_B_, where W stands for fitness. ε is shown as the mean of three biological replicates. (**e**) Checkerboard assay to quantify baseline epistasis between plasmid backbones pC194 and pWL17. RN4220 cells carrying both plasmid backbones, pC194 and pWL17, were plated on TSA containing various combinations of IPTG and xylose and grown for 22 hours at 37 °C. (**f**) Quantification of fitness and epistasis for checkerboard assays in (e). (**g**) Distribution of baseline epistasis (ε) among 12 IPTG-xylose combinations in (f). (**h**) To contextualize the epistasis measured in our study, we re-plotted the distribution of epistasis among ∼15 million gene pairs (top) and 600,000 essential gene pairs (bottom) measured in a comprehensive genetic interaction network study in yeast^5^. (**i**) Select checkerboard assays from (c) that were incubated for an additional 24 hours at room temperature. Arrows indicate combinations of IPTG and xylose that yielded few or no viable colonies. (**j**) Checkerboard assays to quantify genetic interactions between five CC genes and *coaA* and *ribF*. (**k**) Quantification of fitness and epistasis for checkerboard assays in (j). (**l**) Checkerboard assays to quantify genetic interactions between two CC genes and five DD genes in Δ*sosA* background. (**m**) Quantification of fitness and epistasis for checkerboard assays in (l).

**Fig. S9.**
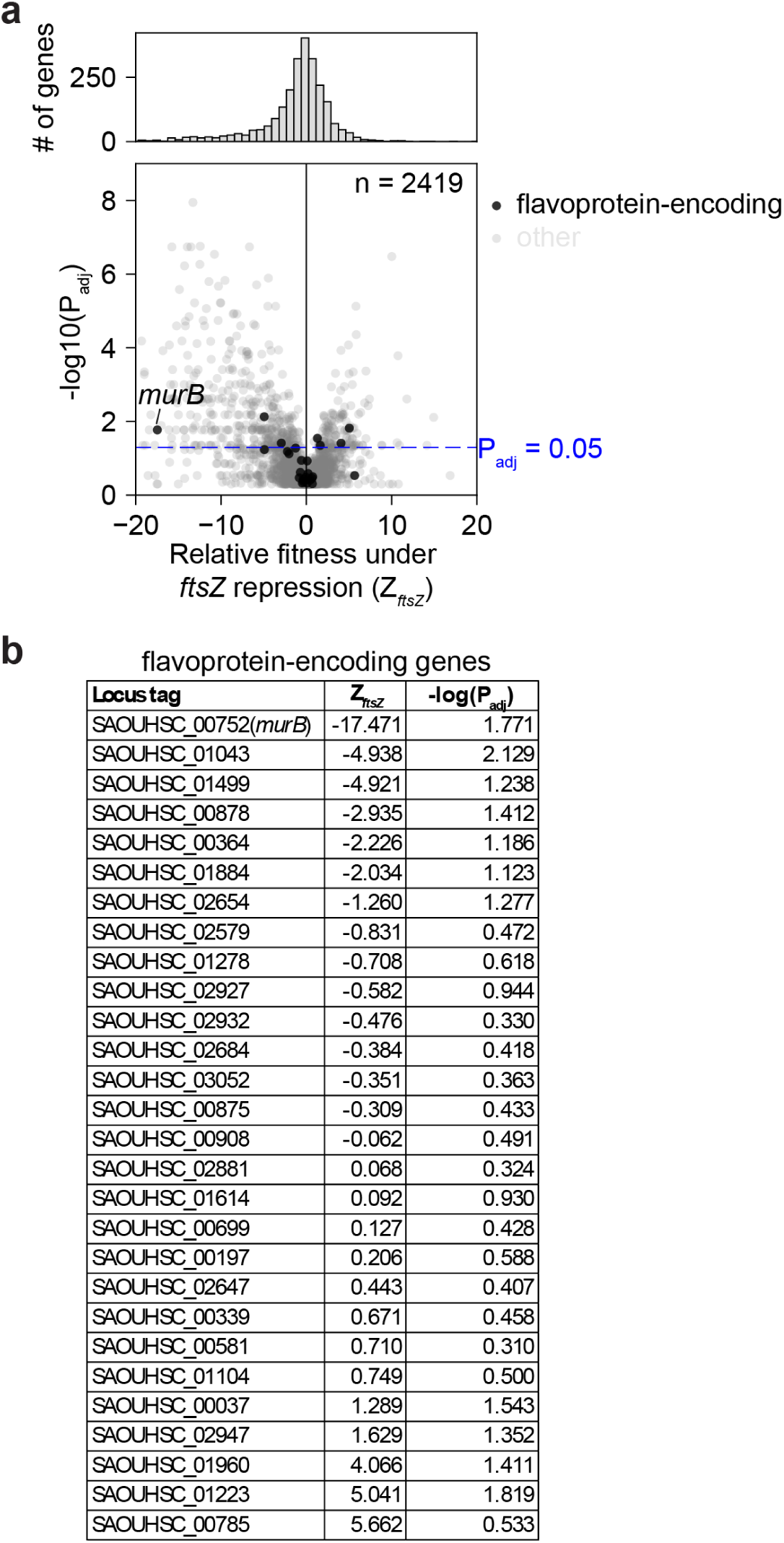
Relative fitness of flavoprotein-encoding genes under mild *ftsZ* repression in *S. aureus* RN4220. (**a**) Volcano plot showing the relative fitness of genes under mild *ftsZ* repression to plain media, Z*_ftsZ_*. Flavoprotein-encoding genes are highlighted in black. (b) Table of Z*_ftsZ_*and -log(P_adj_) for all flavoprotein-encoding genes. List of flavoprotein-encoding genes was obtained from https://www.uniprot.org/uniprotkb?dir=ascend&query=%28organism_id%3A93061%29+AND+%28cc_cofactor_chebi%3A%22CHEBI%3A16238%22%29&sort=gene

**Fig. S10.**
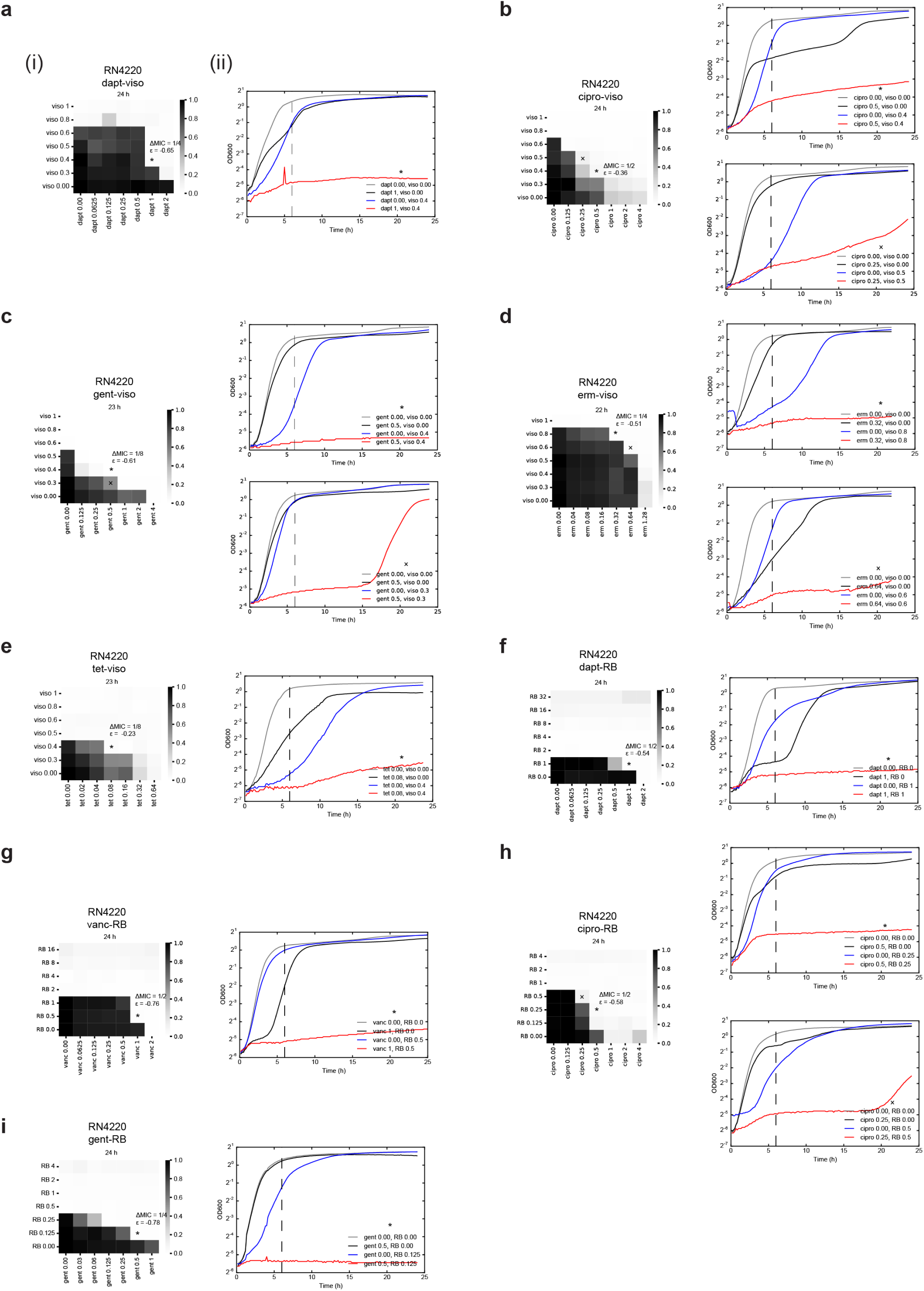
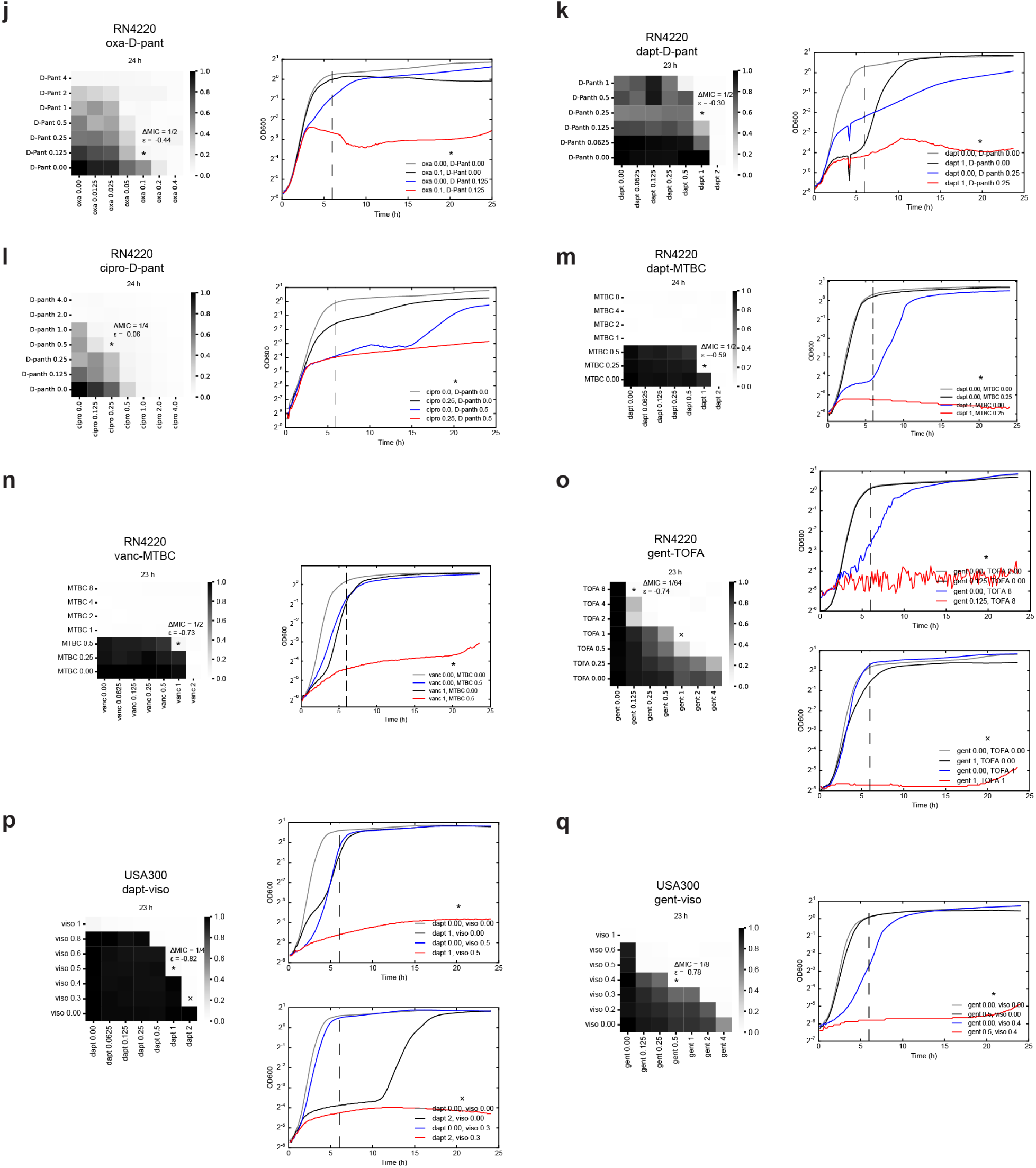
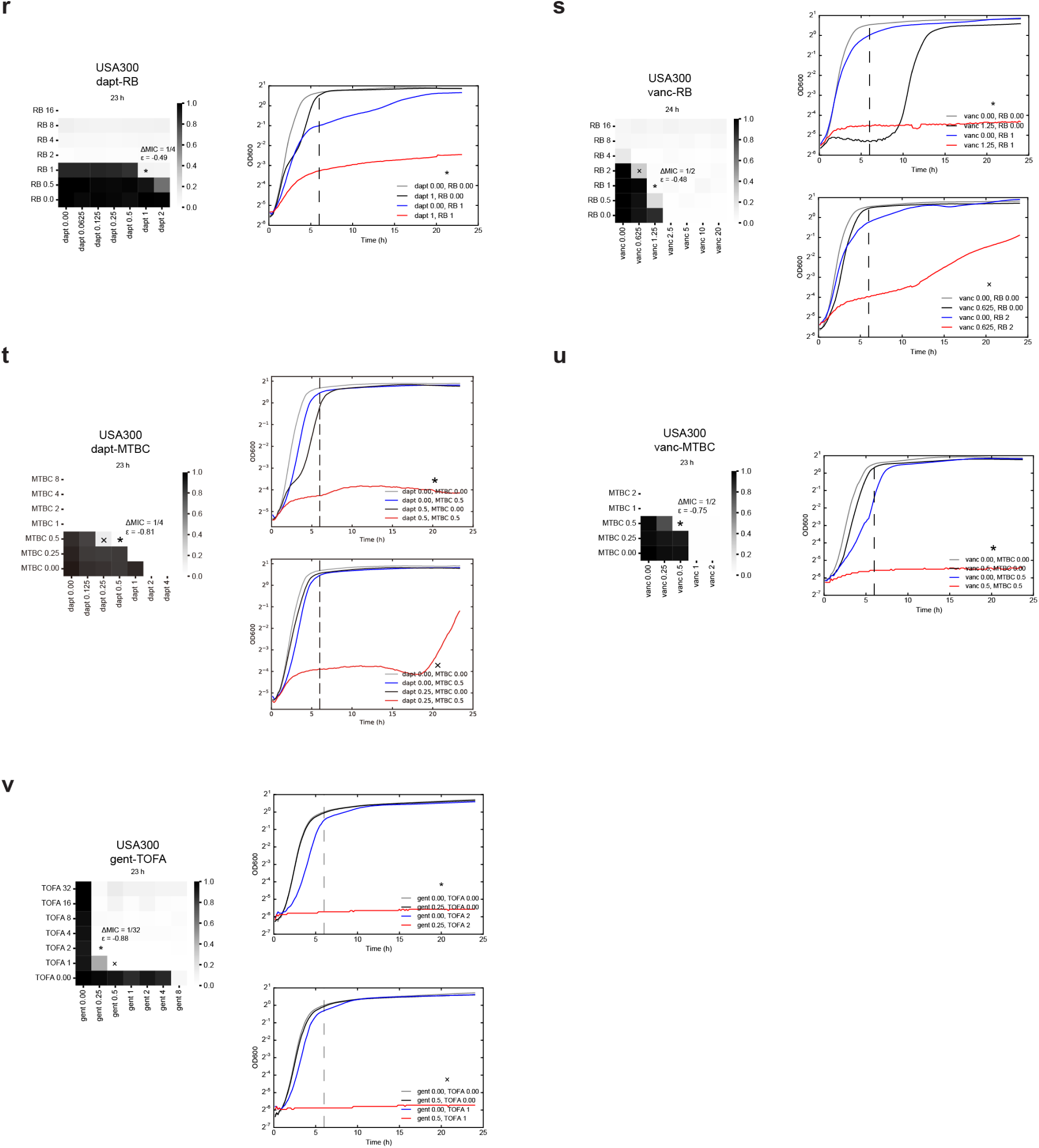
Checkerboard assays for antibiotic-SMI interactions. (**a**) Checkerboard assay of daptomycin-visomitin combination for *S. aureus* RN4220. Panel (i) shows bacterial growth at various concentrations of antibiotic and SMI, normalized to bacterial growth in plain media (ie, 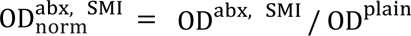) 22-24 hours post inoculation. An asterisk “*” labels the well where the change in MIC (ΔMIC) and epistasis (ε) were calculated (Methods). Panel (ii) shows the four growth curves related to the well labeled with asterisk: plain media (gray), antibiotic alone (black), SMI alone (blue), and both antibiotic and SMI combined (red). Dotted line indicates time = 6 h. Sometimes, a second well with similar growth profiles to “*” was labeled with “x”. The four growth curves related to “x” were also shown. Concentrations of most SMIs and antibiotics are in μg/mL with an exception: the concentration of D-pantothenol is in mg/mL. (**b-o**) Checkerboard assays for other antibiotic-SMI combinations in *S. aureus* RN4220. (**p-v**) Checkerboard assays for antibiotic-SMI combinations in *S. aureus* USA300.

